# Interferon-induced OASL restricts ADAR1 p150 RNA binding and A-to-I RNA editing

**DOI:** 10.64898/2026.10.05.756717

**Authors:** Stephanie Leon Quinonez, Amit Gupta, Soowon Kang, Zhongyi Jiang, Cheng Chih Hsu, Jianrong Li, Feixiong Cheng, Jae U. Jung

## Abstract

The interferon (IFN)-inducible RNA editor ADAR1 p150 catalyzes adenosine-to-inosine (A-to-I) editing of host and viral double-stranded RNAs, limiting aberrant innate immune sensing and regulating gene expression. How ADAR1 p150 activity is regulated during host–virus interactions remains poorly understood. Here, we identify the IFN-stimulated gene OASL as an inhibitor of ADAR1 p150. OASL engages p150 through two distinct interactions: its N-terminal OAS domain associates with the RNA-binding domains in an RNA-dependent manner, whereas its C-terminal ubiquitin-like domains bind the Zα domain independently of RNA. Despite their distinct canonical functions, the OAS and ubiquitin-like domains converge to limit ADAR1 p150 RNA binding and suppress A-to-I editing across the transcriptome. OASL also reverses p150-mediated suppression of antiviral and innate immune gene expression and counteracts p150-mediated enhancement of measles virus and vesicular stomatitis virus replication. Thus, OASL restrains ADAR1 p150, linking interferon signaling to the control of RNA editing and antiviral immunity.

## INTRODUCTION

Activation of the innate immune system via interferon (IFN) signaling is a frontline defense against intracellular pathogens.^1–5^ During RNA virus infection, pattern recognition receptors (PRRs) can sense viral double-stranded RNA (dsRNA), triggering the expression of IFN-stimulated genes (ISGs) and initiating rapid immune responses. To avoid mistaken PRR activation by self-dsRNA, host cells use post-transcriptional modifications.^6,7^ Adenosine-to-Inosine (A-to-I) RNA base editing is the most prevalent RNA modification in mammals catalyzed by the highly conserved Adenosine Deaminase Acting on RNA (ADAR) enzyme family.^8^ This A-to-I editing regulates and diversifies the transcriptome, and creates a wobble-like base pairing that prevents self-dsRNA from activating an aberrant IFN response.^9^ Among ADARs, ADAR1 is the most widely expressed across tissues, while ADAR2 and ADAR3 are neuronally enriched. ADAR1 catalyzes hydrolytic deamination at the C6 position of adenosine to inosine (A-to-I), interpreted as guanosine by the translational machinery, primarily in inverted *Alu* and *SINE* repeats within the 3’ untranslated regions (3’UTRs) of long non-coding dsRNA.^10^

ADAR1 exists in two isoforms: the constitutively expressed nuclear p110 isoform and the IFN-inducible cytosolic p150 isoform.^11^ p150 is upregulated via STAT2-IRF9 signaling, enhanced by STAT1, and is distinguished from p110 by its Zα domain, which binds Z-form RNA and contains a nuclear export signal.^12^ Both isoforms share a Zβ domain, three dsRNA-binding domains (RBD1-3), and a deaminase catalytic domain. While p110 and p150 share many RNA substrates, only p150 and ZBP1 uniquely contain the Zα domain able to bind Z-RNA. ADAR1 is essential for survival as *ADAR1* knockout causes lethality in mice due to aberrant type I IFN induction and mutations in ADAR1 lead to interferonopathies like the Aicardi-Goutières syndrome.^7,9^ Notably, A-to-I editing by ADAR1 in 3’UTRs influences transcript stability and gene expression.^13^

While ADAR1 is essential for maintaining endogenous RNA homeostasis, its dysregulation contributes to tumorigenesis and viral proliferation. Aberrant editing has been found in many cancer types, where AZIN1 (antizyme inhibitor 1) serine (S) to glycine (G) substitution at residue 367 has a much stronger tumorigenic effect than its wild-type (WT) form with predicted worse prognosis in cancer patients.^14,15^ The role of ADAR1 in virus infection is context dependent, acting as either pro- or antiviral based on the virus and host interaction.^16,17^ For many RNA viruses, including hepatitis delta virus (HDV), vesicular stomatitis virus (VSV), and measles virus (MeV), ADAR1 promotes replication by two mechanisms: RNA editing to suppress activation of cytosolic RNA sensors and protein-protein interactions to block RNA sensor function.^15^ Isoform-specific functions have also been observed, where nuclear p110 can restrict measles virus, whereas cytoplasmic p150 facilitates its replication by inhibiting PKR activation and IRF3 signaling.^17^

Several regulatory mechanisms of A-to-I editing have been well characterized for the constitutive ADAR1 p110 isoform under steady-state conditions.^18–20^ Editing efficiency depends on RNA secondary structure, subcellular localization, and homodimerization.^21,22^ Post-transcriptionally, SUMOylation of ADAR1 at lysine 418 within the nucleus can alter editing activity.^19^ In addition, multiple RNA-binding proteins influence editing by modulating ADAR1 expression, protein–protein interactions, or binding to repetitive *Alu* elements.^20^ In contrast, regulation of the IFN-inducible ADAR1 p150 isoform is far less understood, despite its dominant role in preventing aberrant IFN responses during viral infection. Unlike p110, p150 is primarily cytoplasmic and induced by IFN signaling through an alternative promoter, positioning it at the frontline of host–virus interactions. IFN also promotes β-TRCP–mediated ubiquitination and degradation of p110, thereby shifting the balance toward p150 activity during infection.^18^ Functionally, p150 facilitates replication of several viruses by editing or masking viral and endogenous double-stranded RNAs to evade innate immune sensing.^23^ Yet unlike the p110 isoform, the molecular mechanisms that fine-tune p150-mediated A-to-I editing in the context of IFN responses, and their consequences for viral replication, remain largely elusive.^15,24^

The 2’-5’ Oligoadenylate Synthetase-Like (OASL) is an ISG product within the OAS family, traditionally activated with RNase L to degrade viral RNA.^25^ Unlike other OAS proteins, the N-terminal OASL lacks the catalytic triad but retains a functional double-stranded RNA-binding domain.^25,26^ In addition, OASL harbors unique C-terminal Ubiquitin-like domains (UBLs) that mimic polyubiquitination and potentiate activation of the RNA sensor RIG-I.^27^ OASL is rapidly induced upon virus infection and has been characterized as an antiviral factor.^28^ We recently demonstrated that OASL undergoes liquid-liquid phase separation and interacts with ZBP1 and RIPK3, a process essential for virus-induced necroptosis.^29^ These unique structural and functional properties position OASL as more than a canonical ISG effector, raising the possibility that it may directly regulate other RNA-modifying enzymes such as ADAR1 during IFN responses.

In this study, we demonstrate that OASL interacts with ADAR1 p150 primarily during viral infection. Yeast two-hybrid (Y2H) and RNA immunoprecipitation reveal that OASL engages both the Z-form and B-form RNA binding domains of p150, thereby blocking its association with target RNAs. RNA-seq and DNA sequencing confirm that OASL reduces the number, frequency, and distribution of A-to-I editing events in genes linked to stress response and innate immunity, reshaping global gene expression and antiviral signaling. Notably, OASL antagonizes p150-mediated enhancement of MeV and VSV replication. Our findings identify OASL as the first OAS family ISG to directly regulate ADAR1 p150 editing activity, uncovering IFN-dependent mechanism that constrains A-to-I editing and highlighting OASL as a potential therapeutic modulator in both antiviral defense and interferonopathies.

## RESULTS

### OASL interacts with p150 predominantly during virus infection

Since OASL harbors two unique C-terminal Ubiquitin-Like domains (UBLs) not found in other OAS proteins, we sought to identify potential binding partners by performing mass spectrometry analysis of OASL UBL pulldown complexes (**Table S1**) using glutathione S-transferase (GST)-UBL pulldown in HEK293T cells and Y2H screening (**Table S2**) using OASL as bait against a human lymphocyte cDNA library. Both approaches independently identified ADAR1, specifically the p150 isoform, as a high-confidence OASL binding partner (**Figures 1A and 1B**). We validated this interaction in *S. cerevisiae*, where co-expression of full-length OASL and ADAR1 supported growth under triple dropout (3DO) selection, confirming their interaction in yeast (**Figure 1C**). Notably, mammalian cells encode only two Zα-containing proteins, ZBP1 and ADAR1, both IFN-stimulated genes central to innate immunity. ZBP1 contains two Zα domains, Zα1 and Zα2, which share 36% and 0% identity, respectively, with the Zα domain of ADAR1 based on BLAST protein alignment (**Figure 1D**). We previously demonstrated that OASL UBLs interact with the Zα domain of ZBP1 to promote non-canonical virus-induced necroptosis.^29^ This raised the possibility that OASL may similarly target the Zα domain of ADAR1 through its C-terminal UBL domains to regulate ADAR1 activity during IFN responses.

**Figure 1.**
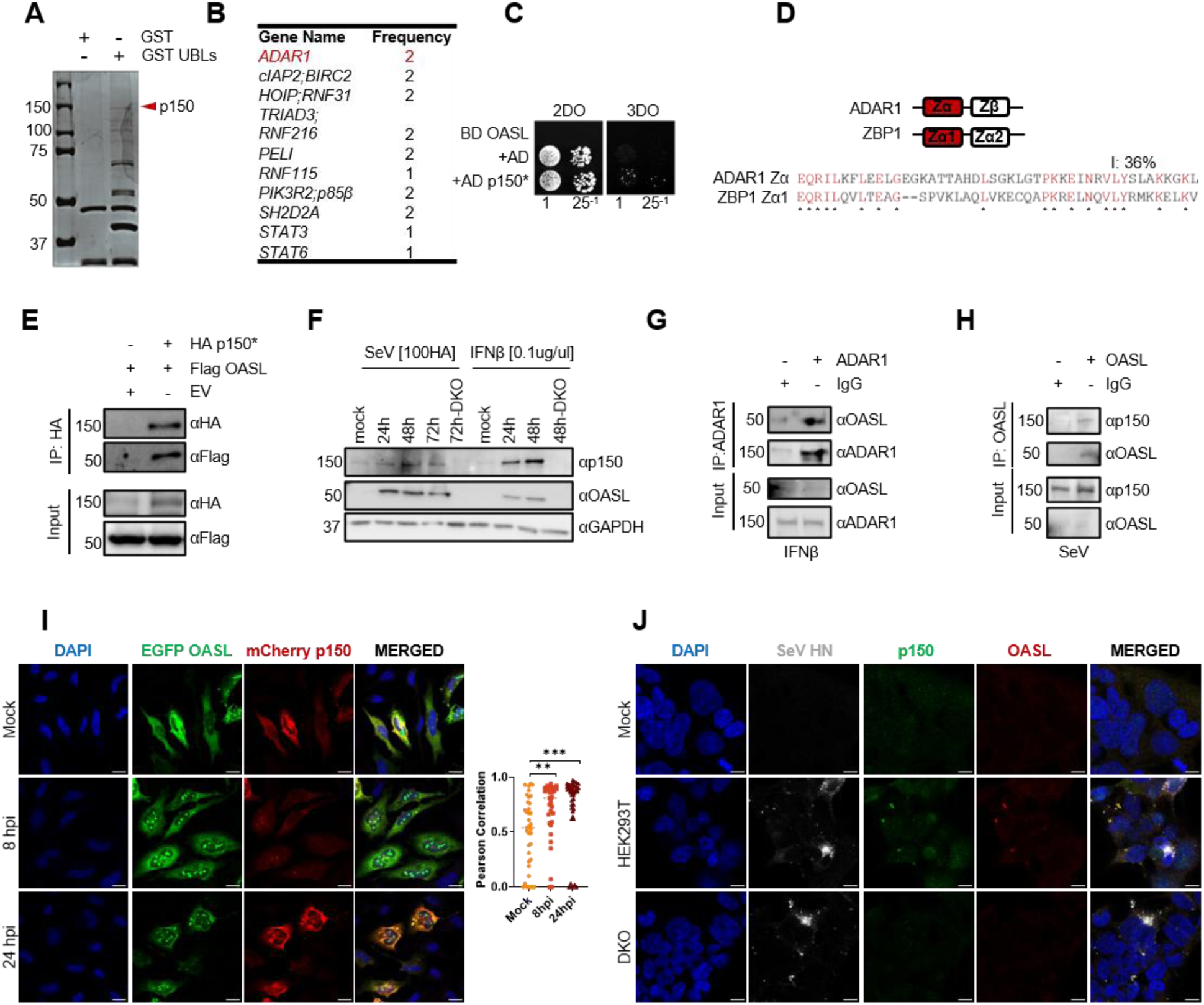
OASL interacts with p150 and co-localizes predominantly during virus infection. (A) Coomassie blue gel used in the mass spectrometry analysis of GST-tagged OASL C-terminus Ubiquitin-like domains (UBLs) pulldown in HEK293T cells. (B) Representation of key yeast two-hybrid (Y2H) assay screening results in *S. cerevisiae* AH109 strain using OASL construct against a human lymphocyte cDNA library. Frequency, number of times prey was identified in replicate screens. (C) Visualization of the Y2H screening of bait OASL with prey ADAR1 p150* construct on an agarose plate.*, M_296_A_297_/L mutation in the start site of p110; 2DO, 2-drop-out agar (-Leu-Trp); 3DO, 3-drop-out agar (-Leu, -Trp, -His); BD, Gal4 binding domain; AD, Gal4 activating domain. Columns indicate 25-fold dilutions from E6 yeast cells. (D) Sequence alignment of human Zα-containing proteins. Analysis performed using BLAST protein alignment tool. ADAR1, Adenosine Deaminase Acting on RNA 1; ZBP1, Z-DNA binding protein 1. (E) Western blot analysis of HA-tagged p150* immunoprecipitation with Flag-tagged OASL using p150* as bait. IP, immunoprecipitation. (F) Immunoblot analysis of endogenous ADAR1 p150 and OASL protein expression in HEK293T cells following Sendai virus (SeV) infection (100 HA, 24 hpi) or IFNβ treatment (0.1 μg/μl, 48 h). DKO, ADAR1/OASL double knockout. (G) Representative immunoblot analysis of endogenous ADAR1 immunoprecipitation from HEK293T cells treated with IFNβ (0.1 μg/mL, 48 h). Immunoprecipitations were performed using an anti-ADAR1 mouse antibody or mouse IgG control. Although the antibody captured both ADAR1 p110 and p150, only p150 is shown. IP, immunoprecipitation. (H) Immunoblot analysis of endogenous OASL immunoprecipitation from HEK293T cells infected with Sendai virus (SeV, 100 HA units, 24 hpi). Immunoprecipitations were performed using an anti-OASL rabbit antibody or rabbit IgG control. ADAR1 p150 was detected using a p150-specific antibody. IP, immunoprecipitation. (I) Fluorescence representation of mCherry-tagged p150 and EGFP-tagged OASL co-localization in HeLa cells at 8h and 24h post-infection (hpi) with Sendai virus (SeV, 100 HA). Left, images taken at 63X and analyzed with ImageJ. Scale bar = 10 µm. Right, Pearson correlation quantification of protein co-localization post-infection. Significance was determined via a two-way ANOVA (*P* < 0.05; \*\**P* < 0.0001). Total number of cells included in analysis (n) for mock, 8 hpi, and 24 hpi are 37, 33, and 41, respectively. (J) Immunofluorescence detection of endogenous OASL (red) and ADAR1 (green) in HEK293T upon SeV (50 HA) infection after 24h. Viral infection was detected by staining for SeV hemagglutinin-neuraminidase (HN), and nuclei were counterstained with DAPI (blue). Images were acquired at 63× magnification using confocal microscopy and analyzed with ImageJ. Scale bar, 10 μm. SeV HN, Sendai virus hemagglutinin-neuraminidase.

We next validated the interaction between OASL and ADAR1 in HEK293T cells by ectopically expressing both proteins. Because ADAR1 is expressed as the constitutive nuclear p110 isoform and the IFN-induced cytosolic p150 isoform, we introduced an M_296_A_297_/L mutation at the p110 start site to selectively express p150 (p150*).^30^ Consistent with our screening results, co-immunoprecipitation confirmed that HA-tagged p150* interacts with Flag-tagged OASL and that OASL UBLs bind specifically to p150 but not p110 (**Figures 1E and S1A**). OASL efficiently associated with both p150 splice variants 1 and 2, despite the truncation in variant 2 within the linker region separating RBD3 from the catalytic deaminase domain (**Figure S1B**). These findings indicate that this linker segment is dispensable for OASL binding and support the use of variant 2 in subsequent functional assays, as it retains the major RNA-binding and catalytic domains required to assess ADAR1 p150-mediated editing and regulation of antiviral signaling.^31^

To determine whether OASL and ADAR1 p150 interact endogenously, we first induced expression of both proteins by treating cells with IFN-β for 48 hours or by infecting cells with Sendai virus (SeV), a potent inducer of type I IFN responses, for 24 hours (**Figure 1F**).^1^ Interestingly, direct IFN-β treatment resulted in higher endogenous expression of ADAR1 p150, whereas SeV infection highly induced OASL expression. These findings are consistent with previous reports showing that rapid IRF3 activation triggered during viral entry promotes early OASL induction whereas ADAR1 expression in human cells is primarily regulated through STAT1–STAT2–IRF9-dependent IFN signaling.^12,28,32^ Immunoprecipitation performed under both conditions demonstrated that endogenous OASL and ADAR1 interact specifically upon enrichment, but not in IgG control samples (**Figures 1G and 1H**).

To assess co-localization under ectopic expression, we infected HeLa cells expressing EGFP-OASL and mCherry-p150 with SeV and observed robust co-localization that increased over time (**Figures 1I, S1C, and S1D**). To investigate co-localization of endogenous OASL and ADAR1 p150, we infected HEK293T and A549 cells and respective knock-out cells with SeV. Using antibodies specific for OASL and the IFN-inducible ADAR1 p150 isoform, we observed co-localization of OASL and ADAR1 p150 in both HEK293T cells (**Figures 1J and S1E**) and A549 WT cells (**Figures S1F and S1G**), whereas these overlapping signals were absent in the corresponding OASL or ADAR1 knockout (KO) cells. Using a SeV hemagglutinin-neuraminidase (HN) antibody as a marker of SeV infection, we observed that OASL and ADAR1 p150 co-localized specifically in the cytosolic puncta in infected cells but showed little overlap with SeV HN, indicating that their association occurs in virus-induced cytoplasmic structures rather than at sites of viral protein accumulation.

Because OASL and ADAR1 p150 co-localized exclusively within cytoplasmic puncta during infection, these observations are consistent with preferential interaction between OASL and the IFN-inducible cytoplasmic p150 isoform rather than the constitutively expressed, predominantly nuclear p110 isoform. Notably, although ADAR1 knockout HEK293T cells remain viable, deletion of ADAR1 is lethal in A549 cells^6^, precluding analysis of ADAR1-deficient A549 cells under identical conditions. Together, these findings demonstrate that viral infection promotes the spatial association of OASL and ADAR1 p150 within cytoplasmic puncta.

### Dual-mode interaction of OASL with ADAR1 p150

To map the interacting domains between OASL and p150, we conducted Y2H assays using a series of OASL constructs as bait and p150* constructs as prey (**Figures 2A and S2A**). The N-terminal OAS domain (N-OAS) strongly interacted with the ADAR1 RBD, with a weaker association observed with Zαβ (**Figure 2B**). In contrast, the C-terminal UBLs exhibited specific interaction with the Zα domain (**Figures 2C and S2B**). Since both OASL and p150 are RNA-binding proteins, we assessed RNA dependency using RNA-binding deficient p150 mutants: W_195_A (Z-RNA binding–deficient)^33^ and EAA (dsRNA-binding–deficient).^34^ The UBL–Zα interaction was independent of Z-RNA-binding activity, whereas the N-OAS–RBD interaction required RNA (**Figure 2D**). Expression levels of RNA-binding mutants were comparable to wild type (**Figure S2C**). Domain dissection revealed that this RNA-independent interaction is mediated specifically by OASL’s second ubiquitin-like domain (UBL2), but not UBL1, with the ADAR1 RBD serving as a negative control (**Figure 2E and S2D**). Additionally, GST pulldown assays with RNase A treatment reinforced that the UBL–Zα interaction occurs independently of RNA (**Figure S2E**). To determine whether OASL and p150 interact directly, independent of cellular cofactors, we purified OASL and ADAR1 domains via affinity and cation exchange chromatography (**Figures S2F, S2G, and S2H**). In vitro binding assays using proteins purified from *E.coli* showed a strong interaction between the GST-tagged UBLs (37 kDa) and the Zα domain alone (10 kDa, **Figure 2F**). On the other hand, in the absence of target RNA, only a weak interaction was observed between GST-tagged N-OAS (62 kDa) and ADAR1 RBD (38 kDa, **Figures 2G and S2I**). An electrophoretic mobility shift assay (EMSA) further confirmed direct interaction between UBLs and ADAR1 Zα, as UBLs displayed a mobility shift upon incubation with increasing concentrations of ADAR1 Zα, whereas no shift occurred in UBL-only controls (**Figure S2J**). Together, these results demonstrate that OASL engages ADAR1 p150 through two distinct protein–protein interactions: the N-terminal OAS domain binds the RBD in an RNA-dependent manner, while the C-terminal UBLs, specifically UBL2, interact with the Zα domain independently of RNA.

**Figure 2.**
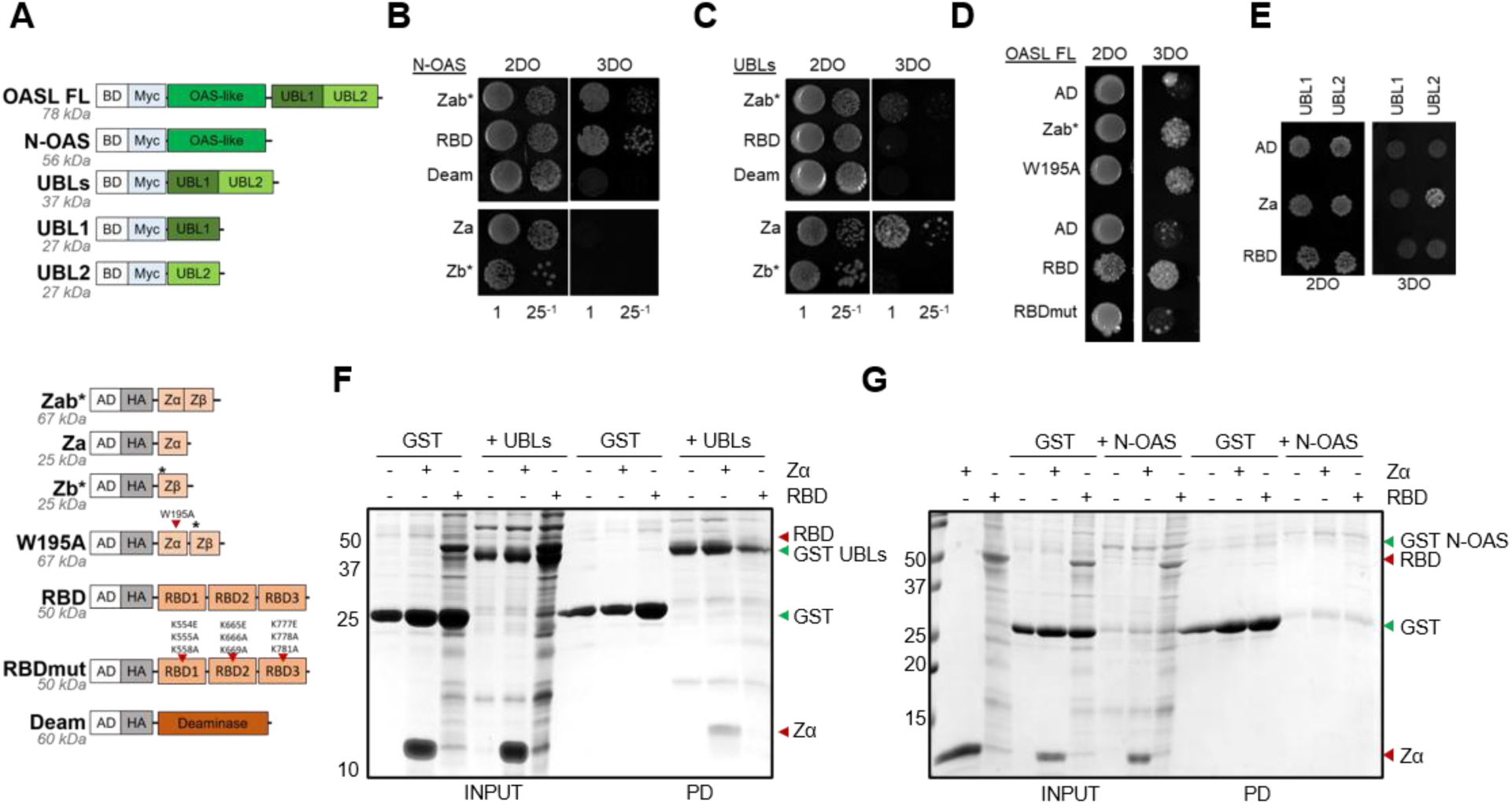
OASL binds to p150 dsRNA-binding domains via its N-terminal OAS (N-OAS) and Ubiquitin-like domains (UBLs) (A) Schematic of constructs used in the yeast two-hybrid (Y2H) domain mapping assay. Top, Myc-tagged OASL constructs in the pGBKT7 bait vector: full-length (FL), N-terminal OAS domain (N-OAS), and the C-terminal Ubiquitin-like domains (UBLs), composed of UBL1 and UBL2. Bottom, HA-tagged p150* constructs in pACT2 prey vector: Zαβ (Zab), Zα (Za), Zβ (Zb), Z-RNA binding mutant Zα mutant (W195A), RNA-binding domains 1–3 (RBD), dsRNA-binding deficient mutant (RBDmut), and the Deaminase domain (Deam). Grey markers indicate predicted molecular weights. BD, binding domain; AD, activating domain. *, M_296_A_297_/L mutation in the start site of p110. (B) Representation of the Y2H assay of *S. cerevisiae* yeast expressing OASL N-OAS domain transformed with respective p150 domains. Columns indicate 25-fold dilutions of initially dropped E6 yeast cells. 2DO, 2 drop-out (-Leu-Trp); 3DO, 3 drop-out (-Leu-Trp-His). (C) Representation of the Y2H assay of *S. cerevisiae* yeast expressing UBLs transformed with ADAR1 Zα and RBD domains. 2DO: 2 drop-out (-Leu-Trp), 3DO: 3 drop-out (-Leu-Trp-His). Columns indicate 25-fold dilutions of initially dropped E6 yeast cells. 2DO, 2 drop-out (-Leu-Trp), 3DO, 3 drop-out (-Leu-Trp-His). (D) Representation of the Y2H assay of *S. cerevisiae* yeast expressing OASL full-length (FL) transformed with empty vector or RNA-binding domain mutants of p150. Yeast colonies initially dropped E6 yeast cells. AD, activating domain; 2DO, 2 drop-out (-Leu-Trp), 3DO, 3 drop-out (-Leu-Trp-His). (E) Representation of the Y2H assay of *S. cerevisiae* yeast expressing each ubiquitin-like domain (UBL) transformed with respective ADAR1 RNA-binding domains. Columns indicate 25-fold dilutions of initially dropped E6 yeast cells. AD, activating domain; 2DO, 2 drop-out (-Leu-Trp), 3DO, 3 drop-out (-Leu-Trp-His). (F and G) Coomassie blue stained SDS gels of GST pulldown assays using purified (F) 6XHis GST UBLs or (G) 6XHis GST N-OAS with 6XHis-tagged ADAR1 Zα or RBD domains. Proteins were purified using cobalt affinity chromatography for purification. Higher–molecular weight bands observed in the gel likely represent non-specific species, as they were consistently detected across all conditions, including the negative controls. The Zα protein was further purified by cation exchange (ÄKTA). Binding performed at 10 μM in 50 mM Tris, pH 7.5, at a 1:1 ratio.

### OASL blocks ADAR1 p150 enzymatic activity by suppressing its binding to target RNAs

Given that OASL binds the ADAR1 p150 RBDs, we investigated whether OASL interferes with RNA binding activity. The p150 dsRBDs, composed of three distinct domains, require duplex secondary structure, with RBD3 being particularly critical for RNA binding and subsequent editing.^35,36^ Unlike ADAR2, which preferentially binds duplexes longer than 11 bp, ADAR1 can edit duplexes as short as 10 bp, relying more on RNA structure than sequence specificity.^35^ To test whether OASL affects p150 RNA binding, we purified the ADAR1 dsRBD1–3 region (RBD) and GST-tagged N-OAS via cobalt affinity chromatography (**Figure S3A**). As target substrates for RNA electrophoretic mobility shift assays (REMSA), we synthesized previously characterized dsRNA substrates known to bind ADAR1 p150: a 38 bp siRNA substrate that supports cytoplasmic ADAR1 p150 binding in an A-to-I editing-independent manner, and a 71 bp RNA substrate derived from the 5-HT2CR pre-mRNA editing region.^37,38^ RBD binding produced a clear mobility shift in both substrates, while N-OAS alone had no effect on RNA migration (**Figures 3A and 3B**). However, increasing concentrations of GST–N-OAS, but not GST alone, caused a reduction in RBD–RNA binding of both target substrates. Coomassie blue staining of SDS-PAGE gels confirmed the presence and relative concentration of each protein in the REMSA reactions. Importantly, N-OAS did not bind the RNA probe directly, indicating it modulates RBD–RNA interactions without competing for RNA.

**Figure 3.**
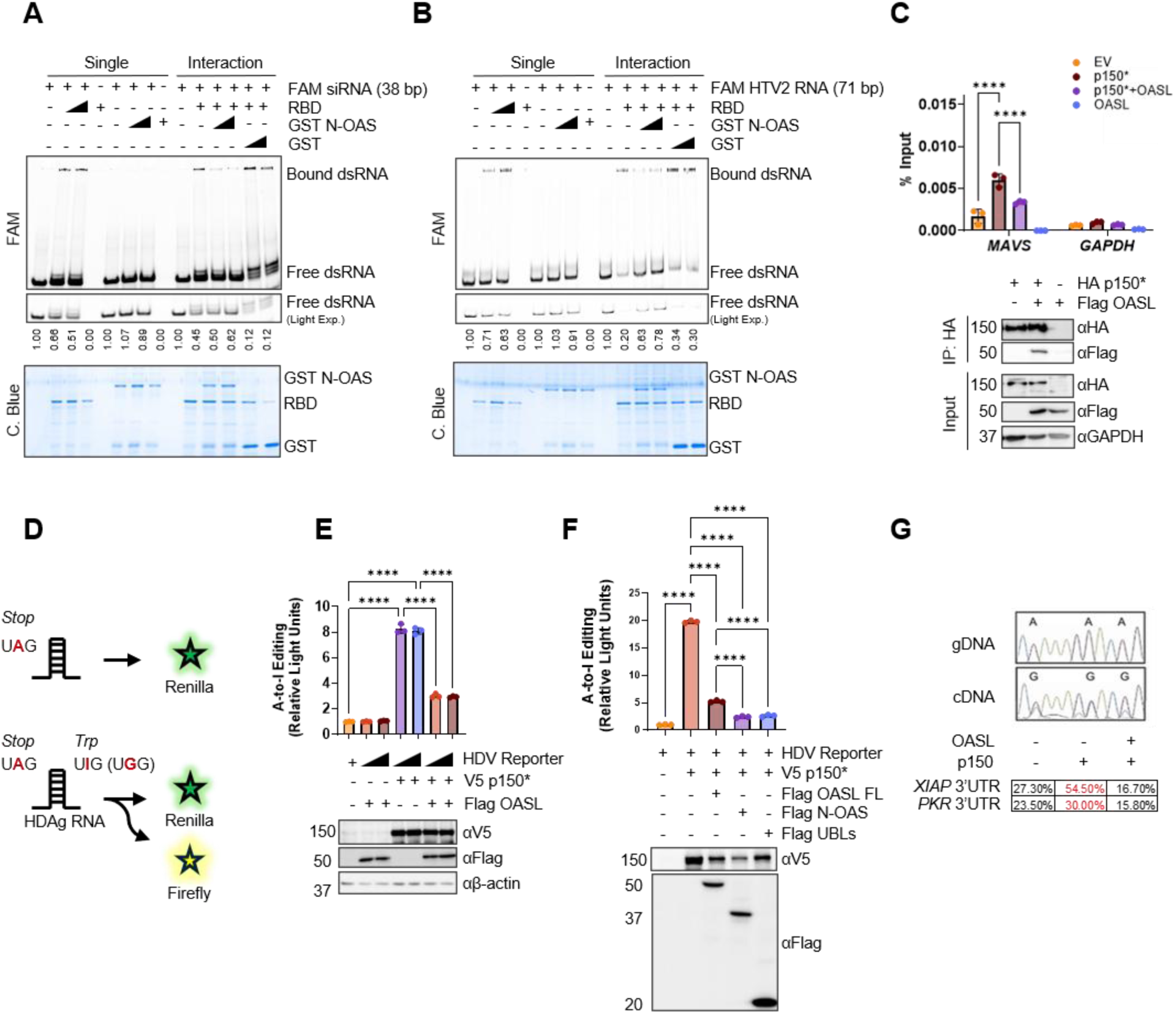
Mechanism of OASL-mediated suppression of A-to-I editing. (A) Fluorescence-based visualization of protein binding to a previously reported ADAR1 p150 substrate consisting of a synthetic 38-bp double-stranded RNA (dsRNA). Top, RNA electrophoretic mobility shift assay (REMSA) performed on a 6% native gel using 5′-FAM-labeled synthetic dsRNA. Numbers represent the ratio of free dsRNA available. Bottom, Coomassie blue staining of SDS-PAGE gels showing protein input and purity. N-OAS, N-terminus OAS domain of OASL. RBD, ADAR1 RNA binding domains. FAM, 5’-fluorescein amidite-label. C. blue, Coomassie blue. (B) Fluorescence-based visualization of protein binding to a previously characterized ADAR1 p150 substrate consisting of a synthetic 71-bp double-stranded RNA (dsRNA) derived from the serotonin 5-HT2C receptor pre-mRNA. Top, RNA electrophoretic mobility shift assay (REMSA) performed on a 6% native gel using 5′-FAM-labeled dsRNA. Numbers represent the ratio of free dsRNA available. Bottom, Coomassie blue staining of SDS-PAGE gels showing protein input and purity. N-OAS, N-terminus OAS domain of OASL. RBD, ADAR1 RNA binding domains. FAM, 5’-fluorescein amidite-label. C. blue, Coomassie blue. (C) RNA immunoprecipitation (IP) assay. Top, Gene expression levels of enriched *MAVS* as %input (2^-dCT^) via RT-qPCR from (C). *GAPDH* was used as negative control for target enrichment. Data shown as mean ± SD from three replicates. Significance was determined via two-way ANOVA (GraphPad Prism), *P* < 0.05; \*\**P* < 0.0001. EV, empty vector; PD, pulldown. Bottom, western blot analysis of pulldown efficiency of RIP using HA-tagged p150* in ADAR1 KO HEK293T cells. (D) Dual-Glo luciferase assay using the Hepatitis D Virus (HDV) reporter. Top, schematic of *Firefly* and *Renilla* luminescence as measurement of A-to-I editing activity. (E) Luminescence results of the Dual-Glo Luciferase assay. Bottom, western blot analysis of expressed constructs in HEK293T cells. Luminescence analysis performed using SoftMax Pro software. Data shown as mean ± SEM (n = 3). Significance was determined via a two-way ANOVA (*P* < 0.05; \*\**P* < 0.0001). (F) Dual-Glo luciferase assay using OASL domains. Top, luminescence results of the Dual-Glo Luciferase assay for measuring A-to-I editing activity. Bottom, western blot analysis of expressed constructs in HEK293T cells. Luminescence analysis performed using SoftMax Pro software. Mean ± SD from two replicates. Significance was determined via a one-way ANOVA (*P* < 0.05; \*\**P* < 0.0001). HDV, Hepatitis D Virus. (G) Target-specific A-to-I editing analyses using Sanger sequencing. Top, diagram depicting A-to-changes in cDNA. Bottom, Sanger sequencing of endogenous editing in the 3′UTR of reported edited targets. Percent editing calculated as G / (G + A) × 100.

We next assessed full-length p150* binding activity to endogenous RNA targets in cells. To avoid interference from the constitutively expressed p110 isoform, we performed RNA immunoprecipitation (RIP) in HEK293T ADAR1 KO cells expressing HA-tagged p150*, followed by RT-qPCR analysis of well-characterized ADAR1 target transcripts (**Figures S3B and S3C**) ^8,39^. Full-length p150* efficiently associated with endogenous transcripts, consistent with recognition of local double-stranded RNA structures.^41^ In contrast, deletion of the Zαβ domain abolished binding to these transcripts, demonstrating that the Zαβ region is required for efficient RNA association by full-length p150*.

We then tested whether full-length OASL could inhibit p150* RNA binding to cellular substrates. Co-expression with OASL significantly reduced ADAR1–RNA association with *MAVS* transcripts without affecting p150* expression or pulldown efficiency (**Figure 3C**). Because ADAR1 enzymatic activity also depends on homodimerization^42^, we tested the effect of OASL on p150 dimerization by co-expressing V5- and HA-tagged p150* with Flag-tagged OASL. Robust p150–p150 interactions were observed and remained unaffected by OASL (**Figure S3D**), indicating that OASL inhibits p150 RNA binding without disrupting its dimerization.

As RNA binding is essential for A-to-I editing, we tested whether OASL impairs ADAR1 p150 catalytic activity. We employed a dual-luciferase reporter containing a 24-nt ADAR1 substrate from the Hepatitis D virus (HDV) amber/W site inserted between Renilla and Firefly luciferase genes.^43,44^ In this system, direct ADAR1-mediated editing converts a stop codon to tryptophan, thereby enabling Firefly expression (**Figure 3D**). We found that editing was primarily mediated by p150 and was significantly reduced by OASL co-expression (**Figure 3E and S3E**). Furthermore, both N-OAS and UBLs domains independently contributed to this inhibition (**Figure 3F**).

To assess whether OASL modulates ADAR1 p150-mediated editing of endogenous substrates, we performed Sanger sequencing of *XIAP* and *PKR* transcripts and quantified A-to-I editing using the G/(G+A) peak height ratio.^45^ Editing levels increased with ADAR1 p150 expression and were significantly reduced upon OASL co-expression in HEK293T wild-type cells (**Figure 3G**). The magnitude of OASL-mediated suppression varied between the two transcripts, consistent with substrate-dependent regulation of ADAR1 editing activity. Notably, low basal levels of editing were detected even in the absence of transfected ADAR1 p150, likely reflecting low endogenous p150 expression in HEK293T cells under steady-state conditions (**Figure 1F**). Together, these results support a model in which OASL inhibits ADAR1 p150 activity by blocking RBD–RNA binding, thereby suppressing A-to-I editing.

### OASL regulates ADAR1 p150-mediated A-to-I RNA editing in mammalian cells

To assess the global impact of OASL on A-to-I editing, we used an ectopic expression system to minimize confounding effects from other IFN-inducible genes activated during viral infection.^46^ Using a lentiviral system, we stably expressed OASL, p150*, both, or empty vector (EV) in HEK293T and A549 cells (**Figure 4A**). HEK293T wild-type (WT) cells were chosen for downstream analysis due to their well-characterized ADAR1 editome.^47^ High-quality total RNAs were extracted and validated (**Figure S4A**), followed by paired-end bulk RNA-seq in triplicate to evaluate the effect of OASL on the global A-to-I editome.

**Figure 4.**
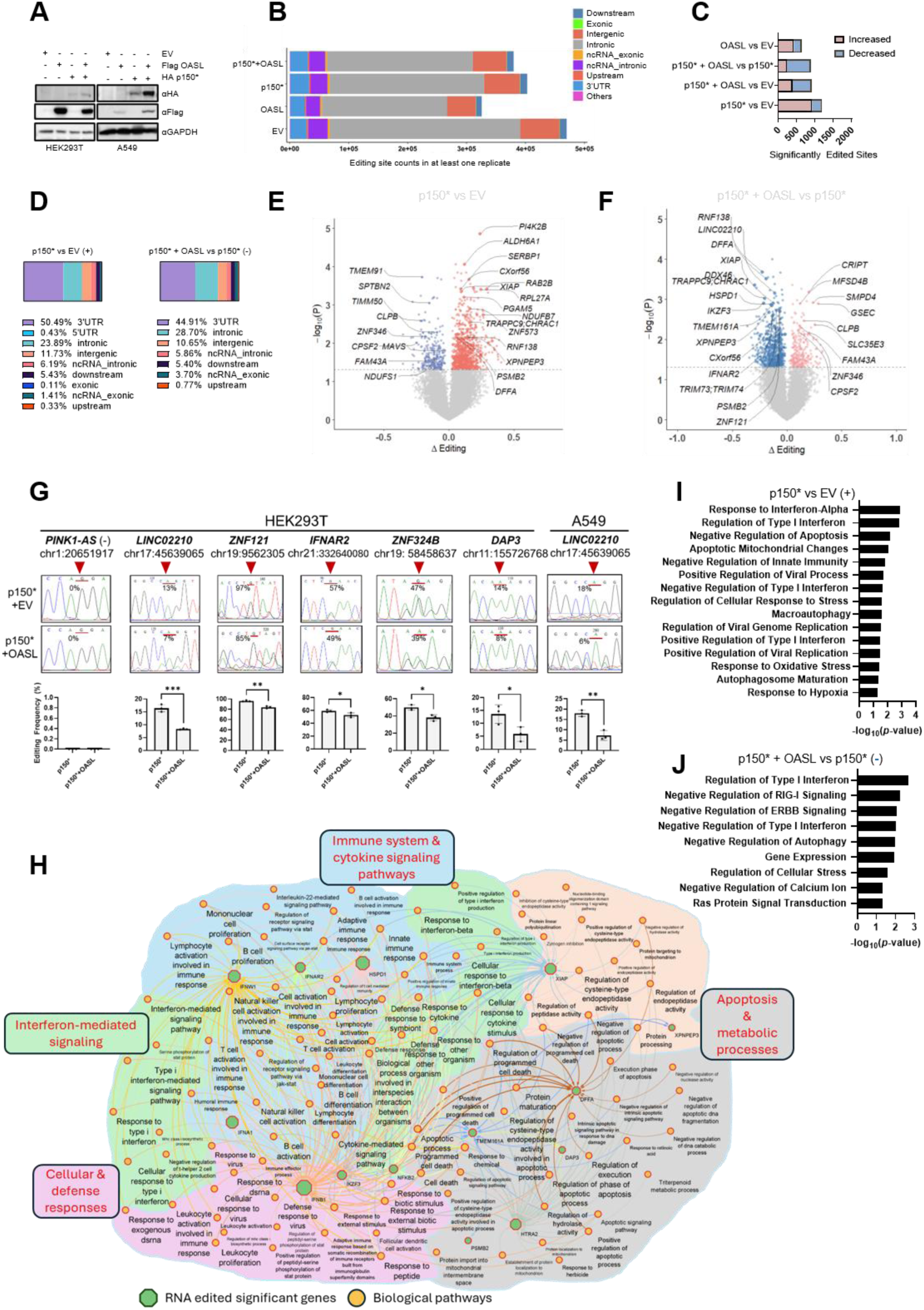
OASL is a potent suppressor of p150-mediated A-to-I editing. (A) Western blot analysis of stably expressing HEK293T and A549 cells. (B) Analysis of the genomic distribution of total editing sites across all samples. Sites included if present in at least one replicate per sample. (C) Bar graph showing the number of significantly increased (left) and decreased (right) RNA editing sites identified in pairwise comparisons of HEK293T transcriptomes. Only editing sites detected in all three biological replicates of each experimental group were included in the analysis. EV, empty vector. (D) Genomic distribution of edited sites. Left, distribution of upregulated sites in the p150* vs EV group. Right, distribution of downregulated editing sites in the p150*+OASL vs p150* group. Percentages indicate distribution across annotated regions. (E and F) Volcano plots of significantly differentially edited sites in HEK293T cells comparing p150* vs. EV (E) and p150*+OASL vs. p150* (F). Red and blue indicate increased and decreased editing, respectively. (G) Sanger validation of selected edited sites in HEK293T and A549 cells. Top, Sanger traces of edited positions with percent editing assessed as G / (G + A)×100. Arrows indicate edited base pair position. Bottom, percent editing analyses compared between samples shown as mean ± SEM from biological replicates. Significance was determined via an unpaired two-tailed Student’s *t*-test: *P* < 0.05; *P < 0.01*. (H) RNA edited selected gene-based pathway analysis representing enriched immune, IFN signaling, virus response, and apoptotic mechanisms from (F). Edges represent interactions between genes and associated pathways, edge colors are specific for the individual genes, label font size for pathways depicts enrichment FDR, and edge width corresponds to number of genes (ngenes) associated to the biological process. (I and J) Enrichr pathway analysis (GO Biological Process 2025) highlighting innate immune-related pathways of significantly edited genes (I) upregulated by p150* in the p150* vs EV group and (J) downregulated by OASL in the p150*+OASL vs EV group.

High-confidence A-to-I editing sites were identified independently in each biological replicate (**Figure 4B**) employing REDItools with stringent filtering for coverage, editing frequency level, and homo-polymeric base substitutions. Genomic annotation revealed that p150* expression increased the proportion of editing events within 3′UTR regions compared to the EV control, consistent with the established preference of ADAR1 for structured dsRNA elements enriched within untranslated regions. We next performed comprehensive annotation of editing sites and further genomic region-specific distribution analysis. Across triplicate datasets, we detected 1192, 882, 909, and 638 significantly altered sites (t-test, p ≤ 0.05) in p150* vs EV, p150*+OASL vs p150*, p150*+OASL vs EV, and OASL vs EV comparisons, respectively (**Figure 4C**). Compared with the EV control, which represents constitutive ADAR1 p110-mediated editing, p150* expression produced the greatest number of editing events, whereas co-expression with OASL reduced both the total sites and frequency of upregulated edits, indicating that OASL suppresses p150*-mediated A-to-I editing.

Genomic distribution analysis revealed that p150*-mediated editing was enriched in 3′UTRs, consistent with the established substrate preference of ADAR1.^39^ Notably, OASL co-expression markedly reduced editing within this genomic region (**Figure 4D**). Consistent with these findings, Volcano plot analysis revealed that p150* expression drove widespread increases in RNA editing relative to the EV control (**Figure 4E**). Conversely, co-expression of OASL broadly attenuated p150*-mediated editing (**Figure 4F**), supporting a global inhibitory effect of OASL on p150-mediated RNA editing. Immune-related transcripts were among the most prominently affected targets in the p150*+OASL versus p150* and p150*+OASL versus EV comparisons, while similar overall editing trends were observed in the p150* versus EV and OASL versus EV datasets (**Figures S4B–S4E**). To validate the RNA-seq findings, we randomly selected ten hyper-edited sites (e.g., IFNAR2) from the p150*+OASL vs p150* and p150* vs EV comparisons, along with one unaffected site (PINK1-AS). Nested PCR and Sanger sequencing confirmed that editing at six of the ten sites was enhanced by p150* and suppressed by OASL, whereas the control site showed no detectable editing (**Figure 4G**). Similar suppression was also observed in A549 cells, indicating that OASL-mediated regulation is not restricted to HEK293T cells.

Gene network analysis of significantly edited RNA targets revealed convergence of many affected transcripts on antiviral and stress-response pathways (**Figure 4H**). Enrichr analyses also showed that genes with altered editing were enriched in innate immunity, RNA metabolism, gene expression, and translation pathways (**Figures 4I and S4F**). Notably, OASL reduced editing in transcripts linked to negative regulation of type I IFN and RIG-I signaling (**Figures 4J and S4G**). Furthermore, OASL decreased not only the number of edited genes but also the editing frequency within individual transcripts (**Figure S4H**). Collectively, these findings establish OASL as a negative regulator of ADAR1 p150–mediated A-to-I RNA editing, thereby constraining editing-driven immune modulation in mammalian cells.

### OASL reverses ADAR1 p150-mediated effects on immune signaling pathways

ADAR1 is known to influence gene expression through protein–protein interactions, RNA binding, and A-to-I editing within coding regions and 3′UTRs.^48^ Given that OASL blocks ADAR1 RNA binding and suppresses editing, we examined whether OASL also alters ADAR1-driven transcriptional changes using the same RNA-seq datasets. Differentially expressed genes (DEGs) were defined by FDR ≤ 0.05 and log₂FC ≥ 0.5. Volcano plots from p150* vs EV (**Figure S5A**), OASL vs EV (**Figure S5B**), p150*+OASL vs EV (**Figure 5A**), and p150*+OASL vs p150* (**Figure 5B**) revealed that p150* alone produced the largest number of DEGs (**Figure 5C**). Both p150* and OASL individually altered global gene expression, but co-expression markedly reduced the number of DEGs, 58% and 43% fewer than p150* and OASL alone, respectively, despite no change in protein levels (**Figure 4A**). As most p150*-regulated DEGs were downregulated, we focused on uniquely upregulated DEGs arising under p150*+OASL co-expression. Venn diagram analyses identified 49 such genes (p ≤ 0.01), enriched for immune signaling and calcium ion homeostasis pathways (Enrichr, p ≤ 0.02, **Figure S5C**).

**Figure 5.**
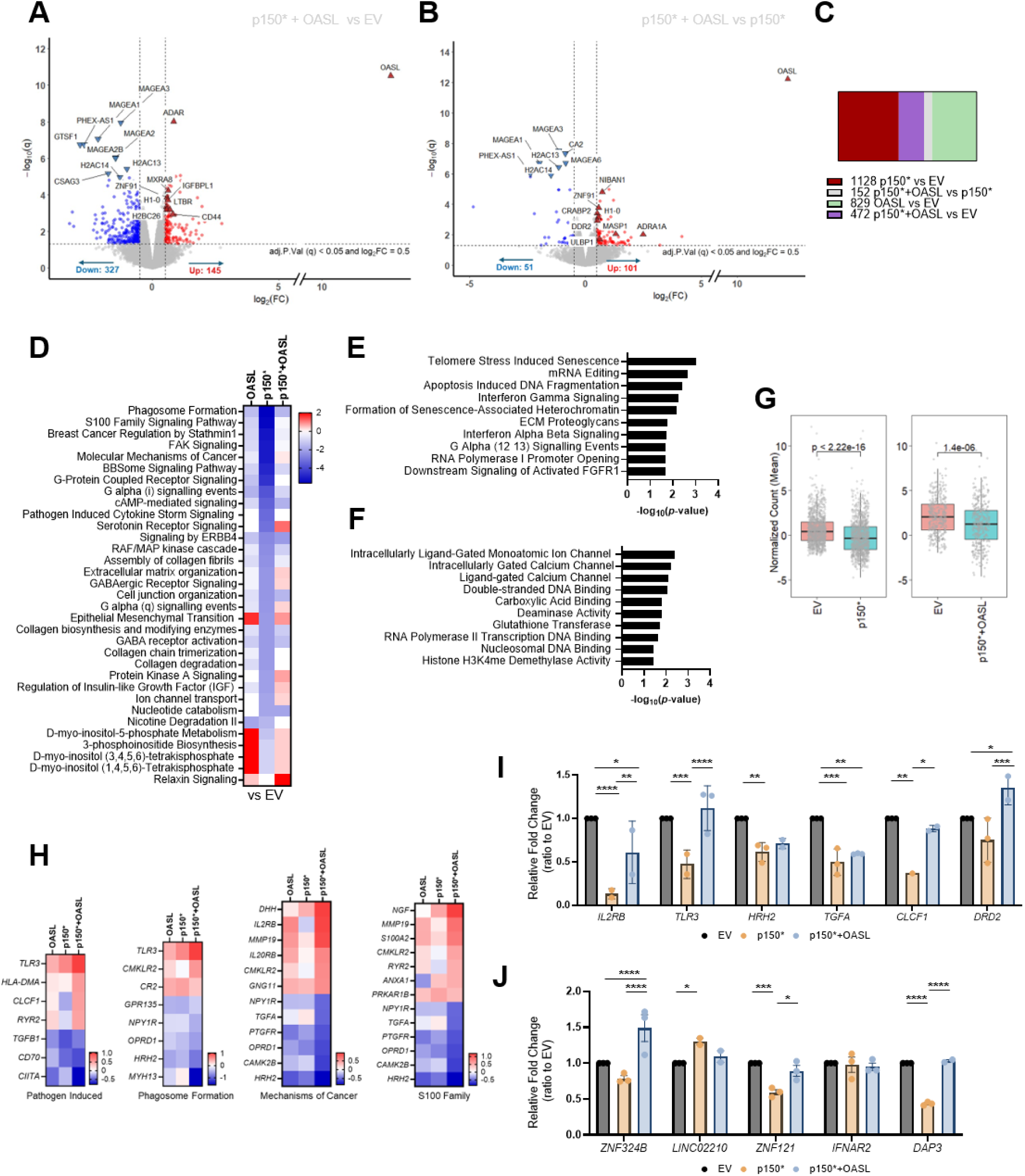
OASL reverts p150-mediated changes in innate immune signaling. (A and B) Volcano plot analyses of differentially expressed genes (DEGs) in (A) p150* vs EV and (B) p150*+OASL vs EV group comparisons in HEK293T cells. Results filtered by FDR < 0.05 and |log₂FC| ≥ 0.5. NS, not significant; FC, fold change; FDR, false discovery rate. (C) Bar graph of the total number of significantly altered genes across groups comparisons. (D) Heatmap of signaling pathways of enriched genes using Ingenuity Pathway Analysis (IPA), filtered by *p* < 0.05 and |Z-score| ≥ 2. Pathways ranked by most suppressed in the p150* vs EV group. EV, empty vector. (E) Enrichr analysis of the Reactome 2024 database showing the top enriched pathways among genes upregulated in the p150* + OASL versus EV comparison. Differentially expressed genes were filtered using p < 0.05 and |log2FC| ≥ 0.5. (F) Enrichr analysis of the Gene Ontology (GO) Molecular Function 2026 database showing the top enriched molecular functions among genes upregulated in the p150* + OASL versus EV comparison. Differentially expressed genes were filtered using p < 0.05 and |log2FC| ≥ 0.5.* (G) Mean normalized expression counts of genes suppressed in the p150* vs EV and p150* + OASL vs EV comparison, shown across the EV, p150, and p150* + OASL conditions. (H) Heatmap of genes from select pathways in (D) showing activation in p150*+OASL vs EV. (I) Gene expression analysis of selected genes from enriched pathways in HEK293T cells via RT-qPCR. (J) Gene expression analysis of significantly edited genes from Figures 4F in HEK293T cells via RT-qPCR.

To determine how OASL-mediated inhibition of ADAR1 p150 affects downstream gene expression, we applied Ingenuity Pathway Analysis (IPA) on pathways most significantly suppressed in the p150* vs EV comparison (p ≤ 0.05, |z-score| ≥ 2). p150* broadly downregulated signaling pathways associated with cellular stress, innate immunity, and cancer (**Figure 5D**). Strikingly, co-expression of OASL reversed much of this suppression, restoring expression of genes involved in pathogen-response, phagosome, S100, and cancer-related pathways. In contrast, OASL alone activated several metabolic pathways that were subsequently suppressed by p150* expression.

To further characterize the underlying biological processes, we analyzed upregulated genes using Enrichr with Reactome and Gene Ontology (GO) Molecular Function datasets for p150* + OASL vs EV (**Figures 5E and 5F**), p150* vs EV (**Figures S5D and S5E**), and OASL vs EV (**Figures S5F and S5G**). These analyses independently supported the IPA findings, demonstrating that co-expression of p150* and OASL upregulated genes associated with IFN signaling, ion channel activity, nucleic acid binding, and apoptotic pathways, despite the distinct transcriptional effects induced by each protein individually.

Upstream regulator prediction analysis (IPA, p ≤ 0.05, |z-score| ≥ 3) further predicted that key pattern recognition receptors suppressed by p150* were less suppressed, or even activated, under p150*+OASL co-expression (**Figure S5H**), consistent with OASL counteracting p150*-driven repression. In addition, we assessed the mean normalized expression of genes suppressed by p150* in both the p150* vs EV and p150* + OASL vs EV conditions and observed that OASL partially restored their expression toward EV levels (**Figure 5G and S5I**). Finally, RT-qPCR validation of six representative genes from immune related pathways independently validated the RNA-seq findings (**Figures 5H and 5I**).

To test whether OASL’s inhibition of A-to-I editing contributes to these transcriptional effects, we examined expression of validated editing targets. Among transcripts suppressed by OASL at the editing level, *ZNF324B*, *LINC02210*, *ZNF121*, and *DAP3* showed significant expression changes in the p150* group, whereas *IFNAR2* did not (**Figure 5J**). However, cross-referencing DEGs with differentially edited genes in the p150* vs EV comparison revealed limited overlap (**Figure S5J**), suggesting that ADAR1 p150 primarily regulates gene expression through editing-independent mechanisms. Together, these findings demonstrate that OASL broadly counteracts p150*-driven transcriptional changes, particularly in pathways governing stress responses, innate immunity, and cancer, thereby reprogramming ADAR1-mediated signaling outcomes.

### OASL suppresses the proviral activity of ADAR1 p150

Our RNA editing and transcriptomic analyses suggested that OASL antagonizes ADAR1 p150-dependent immune suppression. We therefore examined whether this interaction influences viral replication. Stable expression of p150* significantly enhanced MeV Schwarz vaccine strain replication in HEK293T cells by RT-qPCR of MeV *N*^49^, whereas co-expression of OASL reduced viral RNA to near control levels (**Figures S6A, 6A, and S6B**). Similar results were obtained using the pathogenic mCherry-tagged MeV Edmonston strain, where p150* increased infectious virus production and OASL significantly suppressed p150-mediated viral replication, while OASL alone had little effect (**Figures S6C and 6B**).

Because endogenous p110, inducible p150, and other ISGs complicate interpretation in wild-type cells, we next used HEK293T ADAR1/OASL double-knockout rescue cells. Re-expression of p150* restored MeV replication much more strongly than in WT cells, whereas co-expression of OASL consistently suppressed the proviral activity of p150*. OASL alone had little effect (**Figures 6C and S6D**). To determine whether this effect extended beyond MeV, we infected A549 cells with GFP-VSV^15^. As observed for MeV, p150* enhanced VSV replication, whereas OASL reduced p150-mediated viral titers (**Figure 6D**), indicating that OASL broadly antagonizes the proviral function of ADAR1 p150.

**Figure 6.**
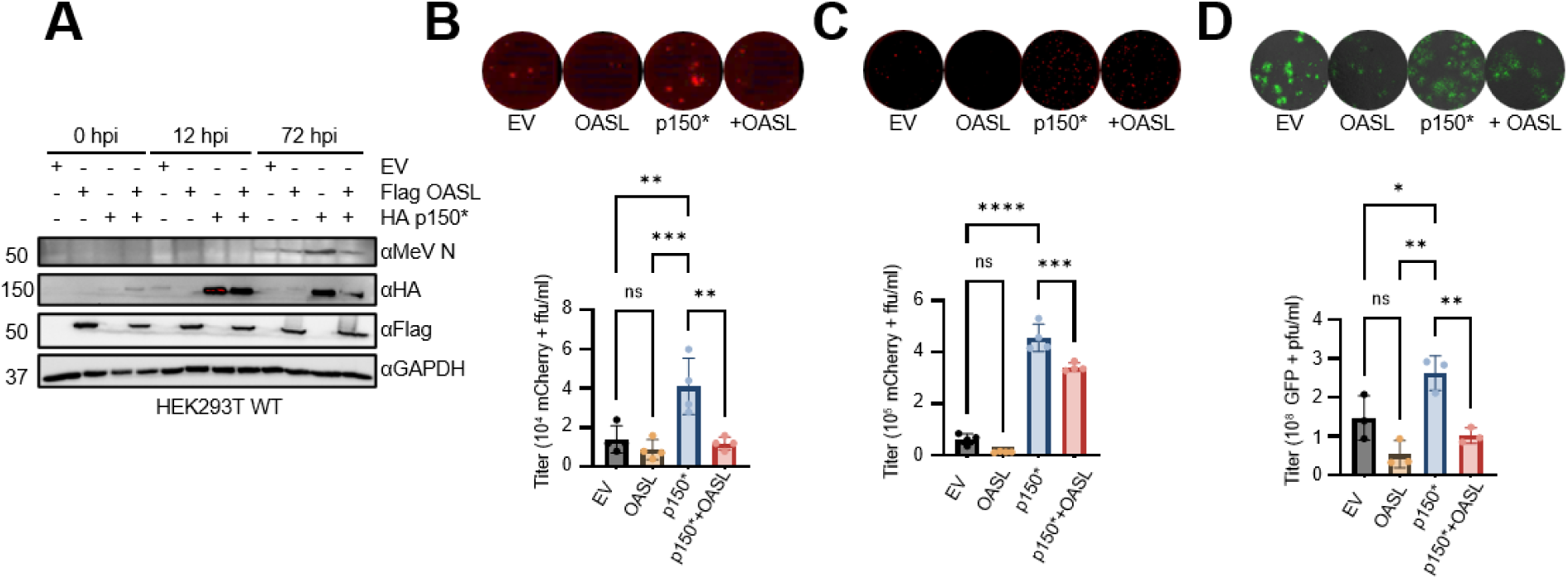
OASL inhibits p150-mediated proliferation of negative-sense RNA viruses. (A) Western blot analysis of MeV N protein in HEK293T cells infected with MeV Schwarz (MOI: 0.1) at indicated hours post-infection (hpi). (B and C) Fluorescence-based focus-forming unit (FFU) assay used to quantify infectious Measles particles. VERO/hSLAM cells were infected with a 1:1000 dilution of mCherry MeV Edmonston virus harvested from A549 WT (C) and HEK293T DKO (D) cells at 3 days post-infection (dpi). Top, representative images from biological triplicates acquired at 5× magnification using an inverted microscope plate scanner (Cleveland Clinic Research Imaging Core). Bottom, quantification of viral titers. Data are presented as mean ± SD (n ≥ 3). Statistical significance was determined by one-way ANOVA: *P < 0.05, ****P < 0.0001. (D) Fluorescence-based plaque-forming unit (FFU) assay used for VSV titration. VERO/hSLAM cells were infected with 10^-7^ dilution of GFP VSV harvested from A549 at 24 hours post-infection (hpi). Top, representative images from biological triplicates acquired at 4× magnification using a Keyence microscope. Bottom, quantification of viral titers. Data are mean ± SD (n=3). Significance determined using a one-way ANOVA: *P* < 0.05; \*\**P* < 0.0001.

We next examined whether the opposing effects of OASL and ADAR1 p150 on viral replication correlated with changes in antiviral cytokine responses. Both MeV and VSV infection induced robust IFN and ISG expression (**Figure S6E**). While expression of p150* reduced antiviral cytokine induction, co-expression of OASL partially restored the expression of type I IFN and IFN-inducible cytokines, with the most pronouced effects observed during MeV infection. We further examined antiviral signaling in MeV-infected HEK293T ADAR1/OASL double-knockout (DKO) rescue cells. Re-expression of p150* increased MDA5 phosphorylation; conversely, co-expression of OASL reduced MDA5 phosphorylation toward EV levels (**Figure S6F**). Consistent with the viral replication phenotype, p150* increased viral mCherry expression, whereas OASL co-expression effectively reduced viral protein accumulation (**Figure S6F**). Together, these results show that OASL-mediated suppression of the proviral activity of ADAR1 p150 is accompanied by changes in antiviral cytokine expression and innate immune signaling during infection.

## DISCUSSION

RNA editing by ADAR1 represents a fundamental layer of post-transcriptional gene regulation that safeguards cells from aberrant activation of innate immunity. As the most prevalent RNA modification in metazoans, A-to-I editing is indispensable for maintaining immune homeostasis, yet its dysregulation drives pathology across diverse settings, from interferonopathies and cancer to viral infection. The IFN-inducible p150 isoform acts as a key suppressor of IFN signaling. Precise regulation of this activity is essential, as dysregulated editing has been linked to interferonopathies, embryonic lethality in mice, Aicardi–Goutières syndrome, lung cancer progression, and enhanced viral replication in humans.^50^ Although cellular control of ADAR1 is evident, most characterized regulators (e.g., DHX9 and DAP3) act on the constitutive nuclear p110 isoform in uninfected cells. ADAR regulators can modulate editing through diverse mechanisms such as limiting protein expression, blocking homodimerization, hijacking RNA substrates, altering localization, remodeling RNA structures, or directly targeting the deaminase domain.^20,45^ Although typically expressed at lower levels than p110, cytosolic ADAR1 p150 is more catalytically active.^51^ These observations suggest that cells must enforce stringent control over p150 activity under IFN conditions. Several ISG pathways, including the MDA5, PKR, and ZBP1 axes, are functionally antagonized by ADAR1 p150 through its RNA-editing and dsRNA-binding activities. However, despite extensive characterization of these reciprocal interactions, direct inhibition of ADAR1 p150 enzymatic or RNA-binding activity by a specific ISG has remained poorly defined. ^51,52^ Here, we identify OASL as an ISG shown to directly interact with ADAR1 p150 and restrict its A-to-I RNA editing activity, revealing a previously unrecognized layer of IFN-mediated regulation of ADAR1.

We identified OASL as a novel suppressor of ADAR1 p150–mediated A-to-I editing by binding both its Zα and dsRNA-binding domains. Our data show that OASL interacts with p150 and restricts its association with RNA substrates without disrupting p150 dimerization, thereby limiting A-to-I editing. Interestingly, despite IFN-inducing hundreds of ISGs, ADAR1 did not exhibit increased editing frequency or site expansion in ISGs compared to non-ISGs in IFN-stimulated HEK293T cells.^46^ Combined with the transcriptomic complexity introduced by viral infection, this observation led us to assess OASL function under non-IFN conditions. In this setting, p150 alone globally increased editing, while co-expression with OASL significantly suppressed editing, including at known ADAR1 targets such as *XIAP*, *MDM2*, and *ARPIN*.^39^ Our data further confirm the high specificity of p150-mediated editing for inverted *Alu* (IR*Alu*) elements, particularly within 3′UTRs and introns in HEK293T cells.^51^ OASL suppressed not only the overall number of editing sites but also editing frequency and regional distribution. The lack of enhanced p150 editing at ISGs may reflect the activity of ISG regulators such as OASL, which dampen p150 activity during IFN responses. Functionally, ADAR1-mediated editing of 3′UTRs in *XIAP* and *MDM2* is linked to repression of gene expression via IR*Alus*, independent of mRNA stability or miRNA targeting.^39^ Although our overexpression system may not fully reflect the endogenous stoichiometry and kinetics of IFN-induced OASL and p150 during viral infection, the physical interaction between endogenous OASL and ADAR1 was readily detected following IFN stimulation and viral infection, confirming the physiological relevance of this regulatory axis. Moreover, OASL consistently counteracted the p150-mediated suppression of immune- and stress-related genes. Together, these findings support a model in which OASL acts as an IFN-inducible regulator of ADAR1 p150, restricting excessive RNA editing while mitigating p150-induced changes in innate immune gene expression.

Mechanistically, our findings indicate that OASL inhibits p150 activity through two distinct modes: RNA-dependent binding of N-OAS to the RBDs and RNA-independent binding of UBL2 to the Zα domain. While the regulatory function of OASL’s catalytically inactive OAS-like domain remains underexplored, RNA-binding proteins often induce conformational changes that expose new interaction surfaces. For p150, conformational rearrangements in the RBD are critical for dimerization and catalytic efficiency,^20,53^ raising the possibility that OASL disrupts RNA engagement by altering RBD conformation. Similarly, OASL UBL2 interacts with the Zα domain of ADAR1, a domain that enhances editing activity in vivo.^33^ UBLs mimic K63-linked polyubiquitin and activate RIG-I via CARD domain binding,^27^ and our results extend their regulatory role to Zα-containing proteins. Notably, OASL binds both known mammalian Zα proteins, ADAR1 and ZBP1, to modulate their immune functions. Whether UBLs inhibit Z-RNA/DNA binding directly or alter Zα conformation remains an open question.

The role of ADAR1 in viral infection is context-dependent, showing either proviral or antiviral effects depending on isoform, virus, and cellular environment.^17,54^ Typically, nuclear p110 is antiviral, while cytoplasmic p150 supports viral replication.^15^ Our data show that OASL antagonizes this proviral activity, restricting p150-mediated enhancement of MeV and VSV replication. Whether OASL restricts p150-mediated viral replication primarily through changes in host RNA editing, viral RNA engagement, or other RNA-binding-dependent functions of ADAR1 remains to be determined. In steady-state conditions, p150-mediated inosine editing masks viral RNA to avoid MDA5 and PKR activation, preventing inflammatory responses, translational shutdown, and cell death.^51^

Our mapping studies also raised several mechanistic questions. Although N-OAS primarily associated with the RNA-binding domains of p150, it also exhibited weaker binding to the Zαβ region. Because direct interaction between N-OAS and the Zα domain was not detected in either yeast two-hybrid or in vitro binding assays, this weak association may instead reflect RNA-dependent interactions involving Z-RNA or RNA-containing complexes. In contrast, UBL2 directly interacted with the Zα domain, but how this interaction suppresses A-to-I editing remains unclear. One possibility is that UBL2 interferes with Z-RNA recognition or alters the conformation of the Zα domain, thereby impairing RNA engagement by p150. Consistent with this model, deletion of the Zαβ domain reduced substrate RNA binding by p150. Additionally, the limited overlap between differentially edited genes and differentially expressed genes indicates that changes in A-to-I editing do not directly predict transcriptional outcomes. Instead, the effects of RNA editing on gene expression are likely context dependent and influenced by the location, abundance, and functional significance of individual editing events.

Our findings support a model in which ADAR1 p150 and OASL perform complementary functions during the interferon response to balance immune homeostasis and antiviral defense. ADAR1 p150 limits aberrant innate immune activation through its RNA-binding and editing activities, whereas OASL restricts p150 RNA engagement and A-to-I editing and counteracts p150-mediated changes in antiviral gene expression. The extent to which OASL restrains p150 during infection is likely influenced by the kinetics and magnitude of IFN-induced OASL expression, its subcellular distribution relative to p150, and viral mechanisms that antagonize IFN signaling. Together, these findings identify OASL as an IFN-induced regulator of ADAR1 p150 that restricts RNA engagement and A-to-I editing while counteracting p150-mediated transcriptional and proviral effects.

In summary, this study identifies OASL as a previously unrecognized ISG regulator of ADAR1 p150, providing new insight into the fine-tuning of A-to-I editing during IFN responses. By binding Zα-containing proteins in human cells, OASL may modulate the activities of ADAR1 and ZBP1 in both viral infection and tumorigenesis. Clinically, ADAR1 is overexpressed in systemic lupus erythematosus, whereas reduced OASL expression has been proposed as a disease-associated biomarker.^55,56^ This imbalance may reflect an IFN-driven dysregulation or loss of OASL’s repressive function on A-to-I editing. Together, our findings establish OASL as an IFN-induced regulator of ADAR1 p150 and reveal a mechanism by which an ISG directly modulates ADAR1 RNA engagement and A-to-I editing. This OASL–ADAR1 regulatory axis provides a previously unrecognized mechanism for controlling p150 activity during the antiviral response.

## Limitations of the study

The effects of OASL on viral replication were examined only in cell culture, and its impact in vivo remains unknown. Global A-to-I editing analyses were performed in limited cancer cell models and under non-infectious conditions, leaving open whether OASL suppresses viral RNA editing during infection. Moreover, the precise residues mediating OASL-ADAR1 interaction and the potential contributions of other OAS family members require further investigation.

## ACKNOWLEDGEMENTS

We thank the Cleveland Clinic Genomics Core for their expert assistance with library preparation and next-generation sequencing. We also thank the Cleveland Clinic Imaging Core, particularly Dr. John Peterson, for their outstanding support and expertise in confocal and inverted microscopy, which were instrumental to this study. We are grateful to Dr. Joseph Luna for generously providing the ADAR1 HEK293T knockout cells, Dr. Sean Whelan for providing the GFP-VSV virus, and Dr. Balaji Manicassamy for providing the A549 OASL knockout cells. This research received no specific grant from any funding agency in the public, commercial, or not-for-profit sectors. We gratefully acknowledge the generous philanthropic support of Sheikha Fatima bint Mubarak.

## AUTHOR CONTRIBUTIONS

Conceptualization, S.L.Q., J.J.; data - S.L.Q.; investigation, S.L.Q., J.J.; visualization, S.L.Q., A.G.; funding acquisition, J.J.; supervision, S.K., J.J.; transcriptomics, A.G.; data analysis, S.L.Q., A.G.; writing, S.L.Q.; review and editing, S.K., Z.J., C.C.H., J.L., J.J., A.G., F.C.

## INCLUSION AND DIVERSITY

We support inclusive, diverse, and equitable conduct of research.

**KEY RESOURCES TABLE**

Table 1

## CONTACT FOR REAGENT AND RESOURCE SHARING

Further information and requests for resources and reagents should be directed to and will be fulfilled by the Lead Contact, Jae U. Jung

## DATA AVAILABILITY

The mass spectrometry proteomics data generated in this study have been deposited in under accession number . The RNA sequencing and RNA editing datasets generated in this study have been deposited in under accession number . All other data supporting the findings of this study are available within the paper, its Supplementary Information, Source Data files, or from the corresponding author upon reasonable request.

---

**Figure S1.**
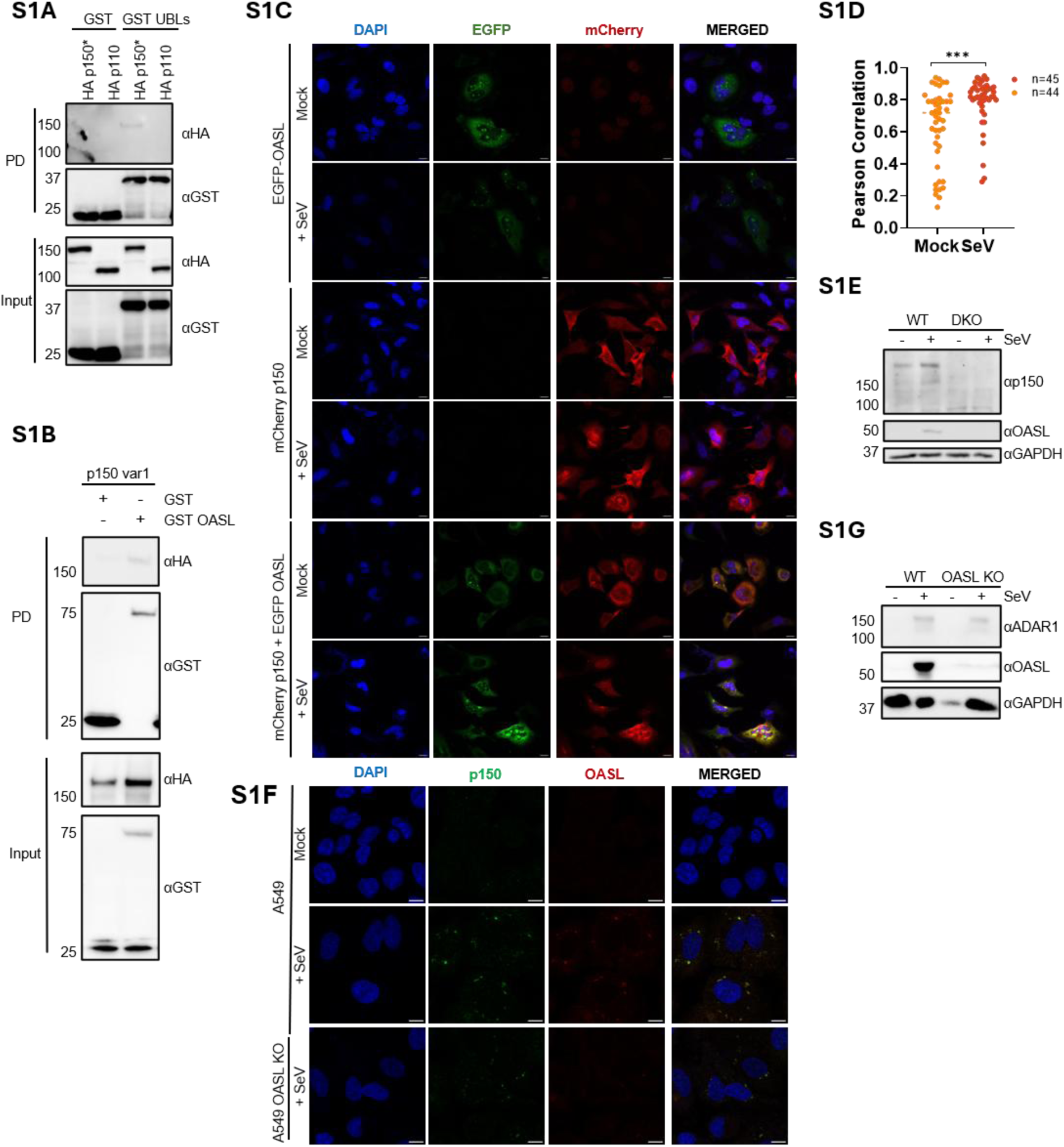
OASL primarily binds p150 variants during viral infection, related to Figure 1. (A) Western blot analysis of GST-tagged Ubiquitin-like domains (UBLs) pulldown with HA-tagged ADAR1 isoforms in HEK293T cells. PD, pulldown; *, mutation in the M_296_A_297_/L start site of p110 within the p150 coding sequence. (B) Western blot analysis of the GST pulldown with ADAR1 p150 variant using OASL as bait. PD, pulldown. PD, pulldown. (C) Fluorescence representation of EGFP-tagged OASL and mCherry-tagged p150 co-localization upon Sendai virus (SeV) infection for 24h via confocal microscopy in HeLa cells. Nuclei stained with DAPI. Scale: 63X. Scale bar= 10 µm. Significance was determined using an unpaired *t*-test: *P* < 0.05; \*\**P* < 0.0001. (D) Pearson correlation from (C) determined via the ImageJ Coloc2 plugin. (E) Western blot analysis of virus-induced ADAR1 and OASL expression in WT and CRISPR-Cas9-generated ADAR1/OASL double knockout (DKO) HEK293T cells after SeV infection. (F) Immunofluorescence detection of endogenous OASL (red) and ADAR1 (green) in A549 cells upon SeV (100 HA) infection after 24h. Cell nuclei were stained with DAPI (blue). Images taken at 63X using a confocal microscope and analyzed with ImageJ. Scale bar = 10 µm. Insets highlight representative infected cells. (G) Western blot analysis of virus-induced ADAR1 and OASL expression in A549 WT and OASL knockout cells following SeV infection.

**Figure S2.**
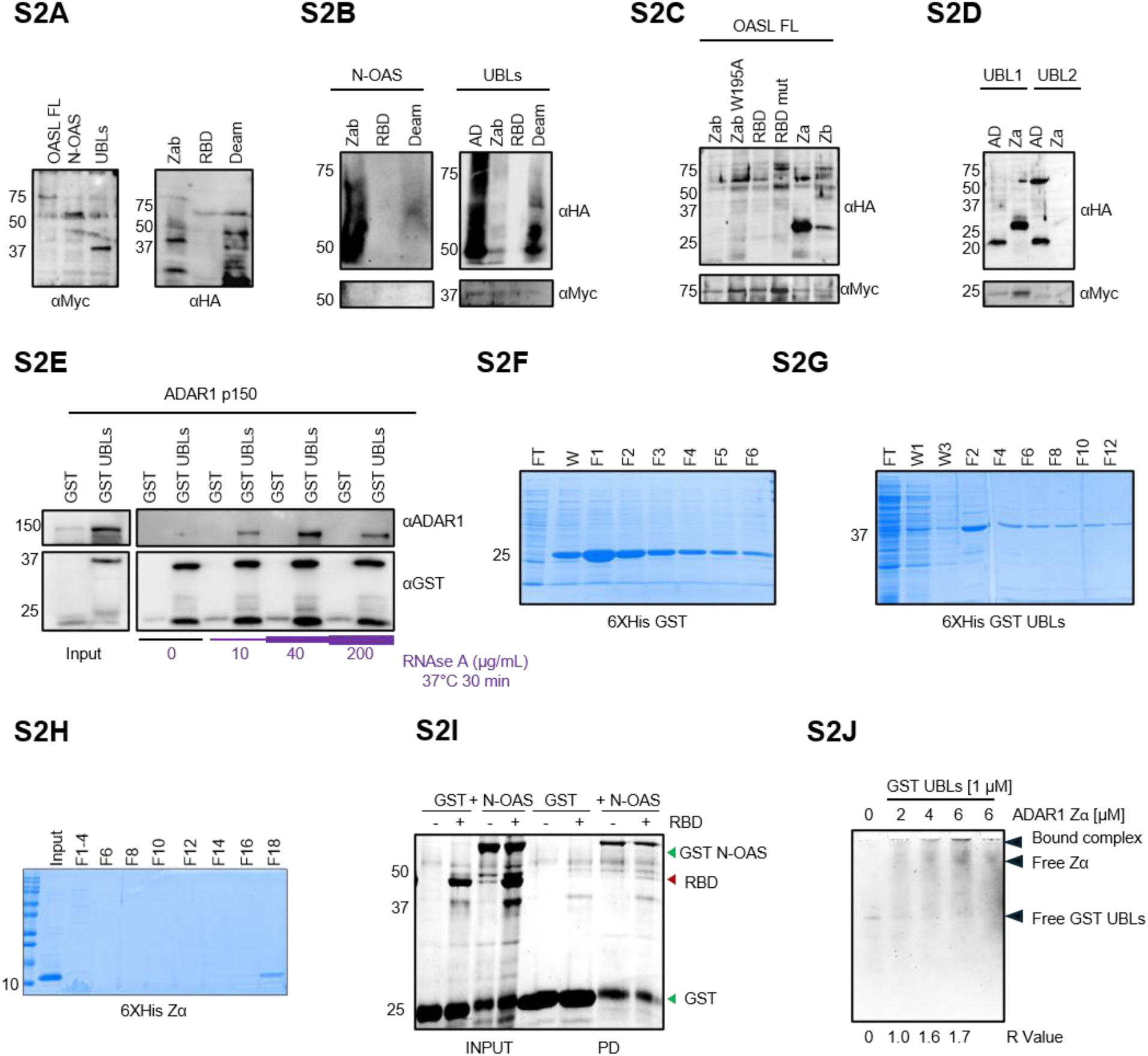
OASL binds to p150 via dual interaction in vitro, related to Figure 2. (A) Western blot analysis of individually expressed constructs in *S. cerevisiae*. Left, pGBKT7-Myc-tagged OASL and indicated domains. Right, pACT2 HA-tagged p150 and indicated domains. (B-D) Western blot analysis of co-expressed constructs in *S. cerevisiae*. (B) Bait pGBKT7 Myc-tagged OASL and pACT2 HA-tagged p150 domain combinations. (C) Bait pGBKT7 Myc-tagged OASL with pACT2 HA-tagged p150* RNA-binding mutants. (D) Bait pGBKT7 Myc-tagged UBL domains with pACT2 HA-tagged p150 Zα domain. (E) Western blot analysis of the GST-tagged OASL UBLs pulldown following increasing RNase A treatment (30 min, 37 °C) in HEK293T cells. (F-G) Coomassie blue stain of the cobalt affinity purification of (G) 6XHis-GST and (H) 6XHis-GST UBLs constructs in Rosetta (DE3) *E. coli* cells. FT, flow-through; W, wash; F, fraction; E, elute. (H) Coomassie blue stain of the cation exchange purification of 6XHis-ADAR1 Zα. A HiTrap 5 mL column on an ÄKTA system was used after cobalt purification. (I) Coomassie blue stained SDS gel (8%) of GST pulldown assay using purified 6XHis GST N-OAS with 6XHis-tagged RBD. Proteins were purified using cobalt affinity chromatography for purification. Higher–molecular weight bands observed in the gel likely represent non-specific species, as they were consistently detected across all conditions, including the negative controls. (J) Coomassie blue stain of the electrophoretic mobility shift assay (EMSA) on a 6% native basic gel using GST-UBLs incubated with increasing concentrations of ADAR1 Zα.

**Figure S3.**
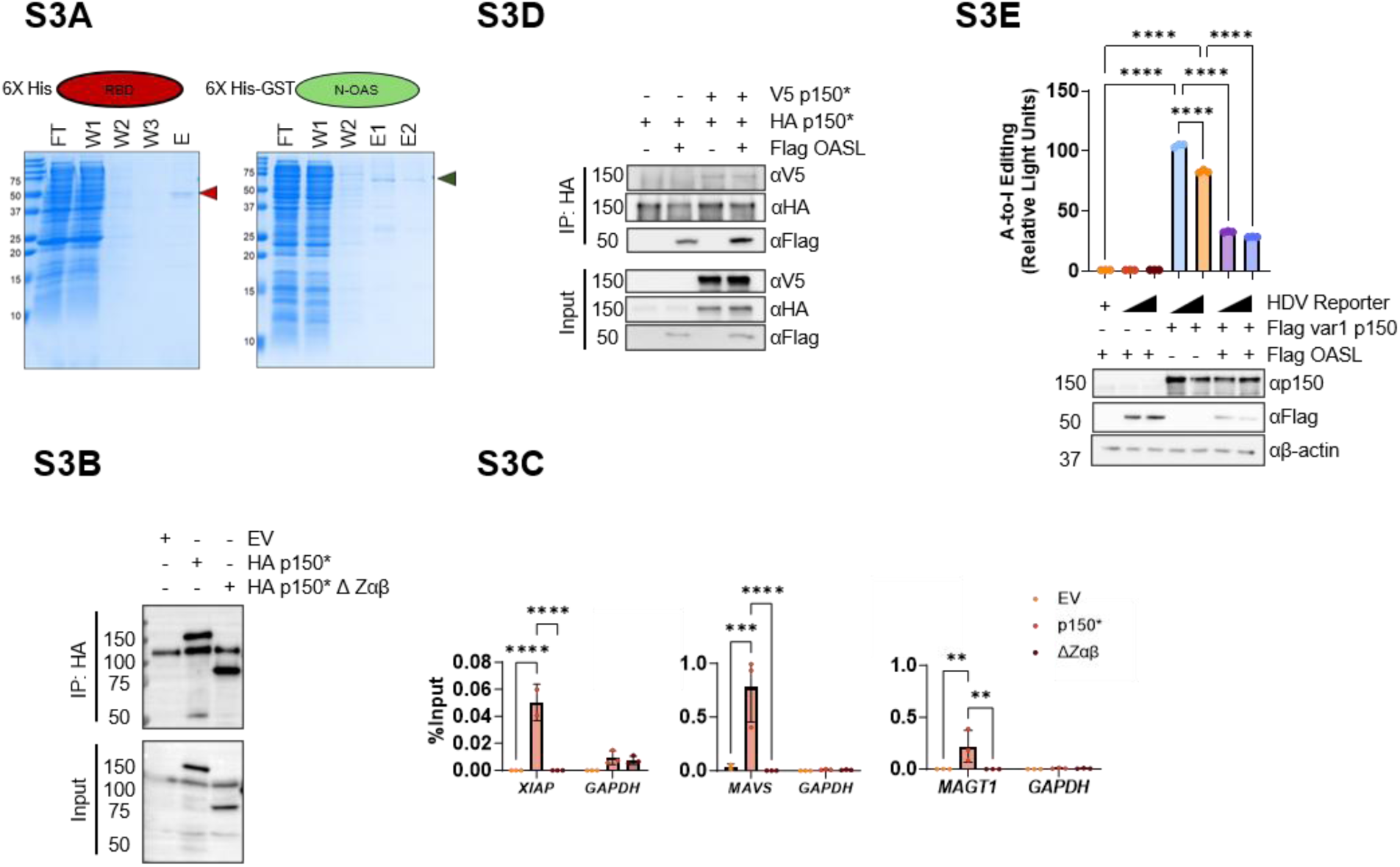
OASL inhibits p150 activity via N-OAS and UBLs domains without altering dimerization, related to Figure 3. (A) Coomassie blue stain of proteins expressed in Rosetta (DE3) *E. coli* and purified via cobalt affinity chromatography. Left, 6XHis ADAR1 RBD construct. Right, 6XHis GST N-OAS construct. Arrows indicate expected molecular weights. FT, flow-through; W, wash; E, elution. (B and C) RNA immunoprecipitation (RNA IP) using full length p150* and respective RNA-binding domain truncations. Right, western blot analysis of RNA IP using HA p150* constructs as bait in ADAR1 KO HEK293T cells. (C) Gene expression analysis of enriched target gene cDNA as % input (2^-dct^) via RT-qPCR. Data shown as mean ± SD (n = 3). Significance was determined via two-way ANOVA. EV, empty vector; PD, pulldown. (D) Western blot analysis of HA/V5-tagged p150* dimerization in the presence of Flag-tagged OASL using ADAR1 KO HEK293T cells. IP, immunoprecipitation; 3XF, 3X Flag. (E) Dual-Glo luciferase assay using multiple variants of the ADAR1 p150 isoform with OASL. Luminescence analysis of the Dual-Glo luciferase assay using the Hepatitis D virus (HDV) editing reporter in the presence of Flag-tagged OASL. A-to-I: adenosine-to-inosine RNA editing. Significance was determined using a one-way ANOVA (*P* < 0.05; \*\**P* < 0.0001).

**Figure S4.**
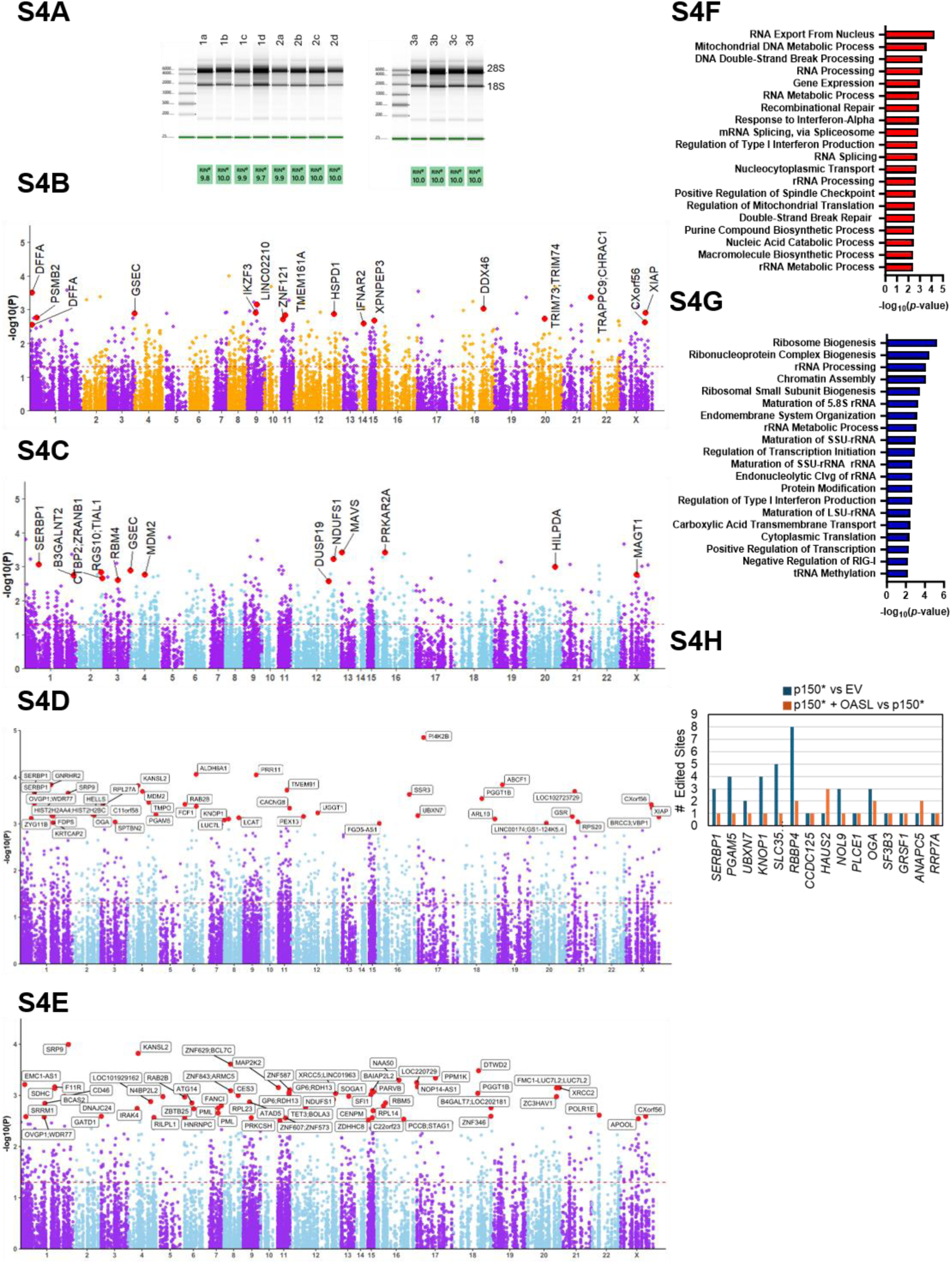
Stable p150 expression enhances A-to-I editing, while OASL co-expression reduces editing, mainly in 3′UTRs of immune genes, related to Figure 4. (A) Gel analysis of RNA quality using a Bioanalyzer of HEK293T stable lines used for deep RNA sequencing in EV (a), OASL OE (b), p150* OE (c), and p150*+OASL (d) groups, each in triplicate (1–3). RIN: RNA Integrity Number. (B) Manhattan plots of significantly differentially edited sites across chromosomes in HEK293T in the p150*+OASL vs p150*. EV, empty vector. (C) Manhattan plots of significantly differentially edited sites across chromosomes in HEK293T in the p150*+OASL vs EV. EV, empty vector. (D) Manhattan plot of significant differentially edited sites in the p150* vs EV group comparison. (E) Manhattan plot of significant differentially edited sites in the OASL vs EV group comparison. (F) Bar graph of top 20 enriched pathways (GO Biological Process 2025) associated with upregulated editing targets in p150* vs EV, identified via Enrichr. (G) Bar graph of top 20 enriched pathways for suppressed editing targets in p150*+OASL vs EV. (H) Graph of editing frequencies of 15 randomly selected significantly edited sites shared between p150* vs EV (blue) and p150*+OASL vs p150* (orange) groups.

**Figure S5.**
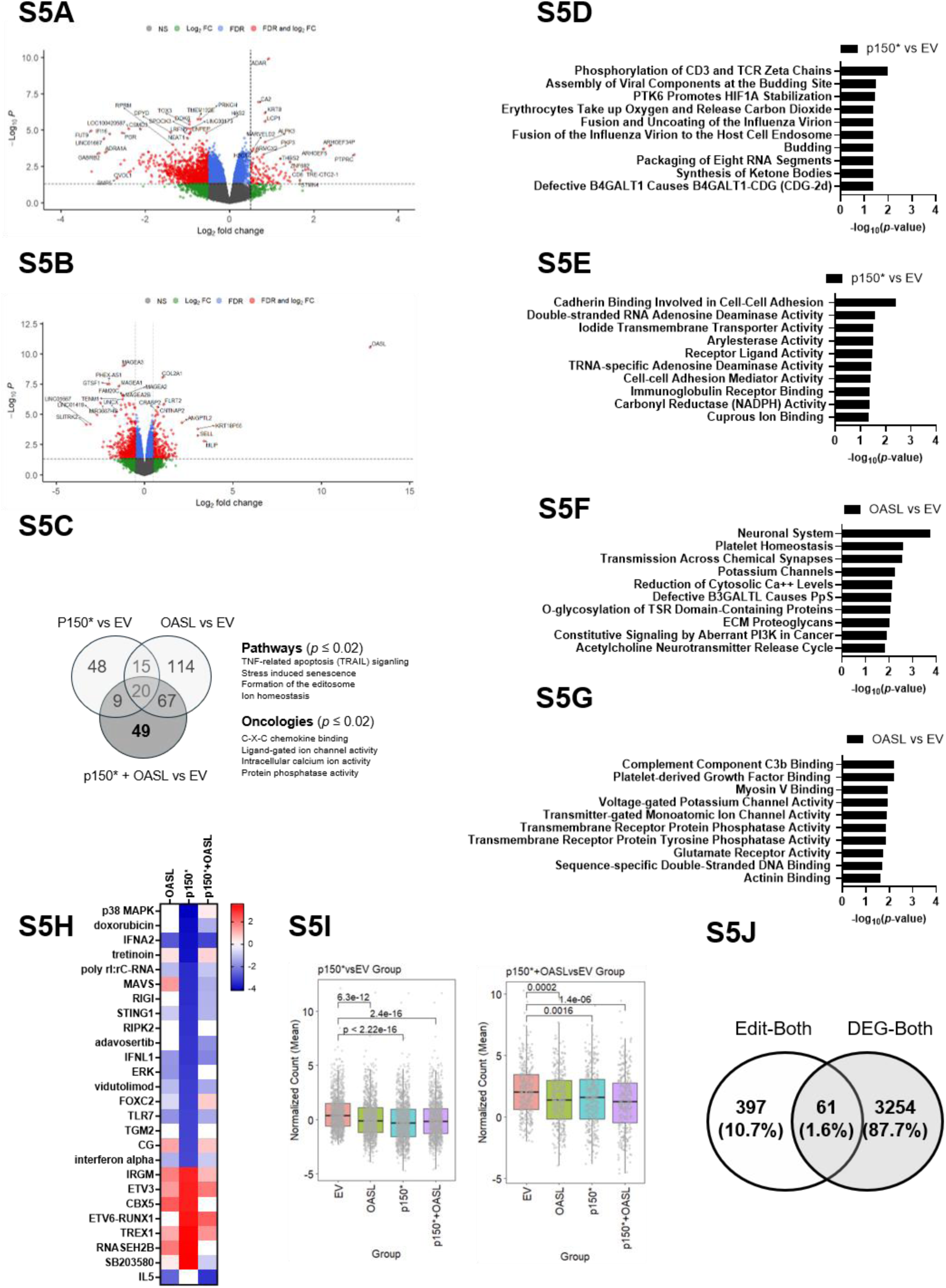
OASL counteracts p150-mediated suppression of predicted upstream regulators and few edited genes show differential expression, related to Figure 5. (A and B) Volcano plots of differentially expressed genes (DEGs). (A) Plot of p150*+OASL vs p150* group comparison. (B) OASL vs EV group comparison. Results filtered by FDR < 0.05 and |log₂FC| ≥ 0.5. NS, not significant; FC, fold change; FDR, false discovery rate. (C) Venn diagram of upregulated DEGs (*p* ≤ 0.01) across group comparisons. Gene overlaps were calculated using the Venny 2.1 online tool. Unique DEGs from the p150* + OASL interaction were subjected to enrichment analysis via Enrichr (Reactome Pathway 2025 and GO Molecular Function 2025). (D and E) Enrichr analysis of the Reactome 2024 database (D) and GO Molecular Function 2026 (E) showing the top enriched pathways among genes upregulated in the p150* versus EV comparison. Differentially expressed genes were filtered using p < 0.05 and |log2FC| ≥ 0.5. (F and G) Enrichr analysis of the Reactome 2024 database (F) and GO Molecular Function 2026 (G) showing the top enriched pathways among genes upregulated in the OASL versus EV comparison. Differentially expressed genes were filtered using p < 0.05 and |log2FC| ≥ 0.5. (H) Heatmap of predicted upstream regulators identified by Ingenuity Pathway Analysis (IPA), filtered by *p* < 0.05 and Z-score ≥ 3. (I) Normalized counts of suppressed genes in each biological replicate from the p150* versus EV (right) and p150*+OASL versus EV (left) comparisons. (J) Venn diagram of gene overlaps between genes with significantly edited sites and differentially expressed genes (DEGs) in the p150*+OAL vs p150* comparison.

**Figure S6.**
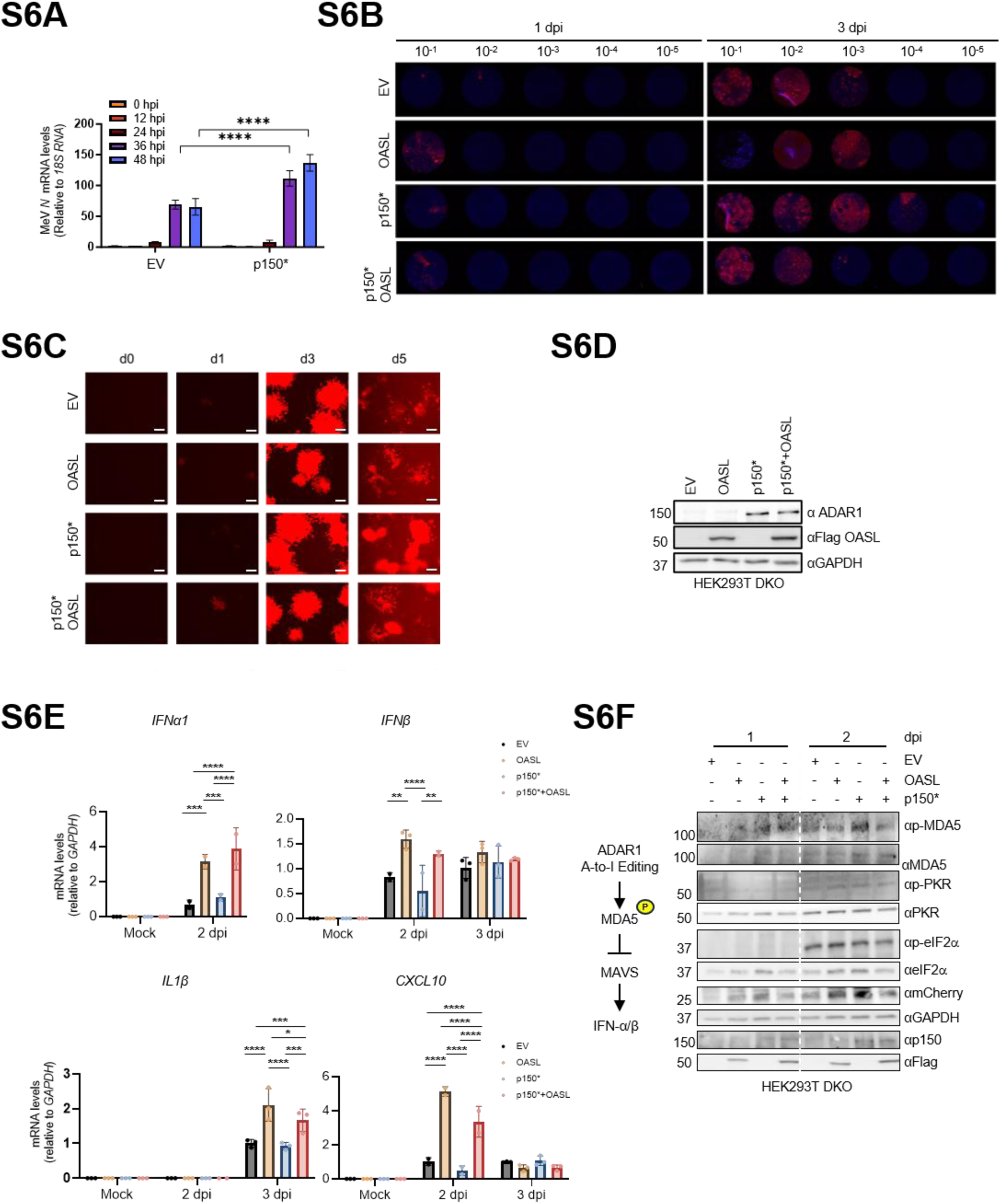
Stable p150 expression promotes measles Edmonston proliferation, while OASL inhibits it, related to Figure 6. (A) Gene expression analysis of Measles virus (MeV) *N* in HEK293T cells stably expressing EV or p150* infected with MeV Schwarz strain (MOI: 0.1) at multiple time points. Data are mean ± SD. Significance was determined via two-way ANOVA: *P* < 0.05; \*\**P* < 0.0001. EV, empty vector. (B) Immunofluorescence analysis of limiting dilution assay of MeV Schwarz propagated in A549 stable lines and titrated in VERO/ hSLAM cells using 10-fold serial dilutions. Detection by MeV N antibody. dpi: days post-infection. (C) Fluorescence analysis of A549 cells infected with mCherry MeV Edmonston strain (MOI: 0.01) at the indicated days post-infection. Supernatants collected for viral titration in Figure 6C. d, day; EV, empty vector. (D) Representative immunoblot analyses corresponding to Figure 6C. (E) Gene expression analyses of cytokines and IFN-stimulated genes (ISGs) in MeV-infected A549 cells. Data are mean ± SD (n = 3). Significance was determined using a two-way ANOVA: *P* < 0.05; \*\**P* < 0.0001. (F) Right, schematic illustrating the signaling pathway suppressed by ADAR1 A-to-I editing. Left, immunoblot analysis of pattern recognition receptor (PRR) signaling pathway activation following infection with mCherry-expressing MeV Edmonston in rescued HEK293T DKO cells. dpi, days post-infection.

## METHOD DETAILS

### Cells

The HEK293T, A549, HeLa cell lines were purchased from ATCC. HEK293T ADAR1 knock-out (KO) cells were generously provided by the laboratory of Dr. Joseph Luna (Case Western Reserve University). A549 OASL KO cells were kindly provided by Sean Whelan (Washington University). HeLa cervical cancer cells, HEK293T human embryonic kidney cells, A549 lung adenocarcinoma cancer cells, and indicated KO cells were cultured in Dulbecco’s Modified Eagle Medium (DMEM) supplemented with 10% heat-inactivated fetal bovine serum (FBS), 1% Penicillin/Streptomycin, and maintained at 37°C in a 5% CO₂ atmosphere. Stably expressing cells in HEK293T and A549 lines were cultured in the DMEM further supplemented with hygromycin (200 ug/ml) and puromycin (1 ug/ml). All cell lines were routinely screened for mycoplasma contamination using the MycoAlert Mycoplasma Detection Kit (Lonza) and confirmed to be free of infection prior to experiments.

### Plasmid constructs

Downstream cloning used the ADAR1 p150 variant 2 template carrying predominantly non-domain mutations (S35P, R100G, K384R, E520D) and the M_296_A_297_/L p110 start site (*) mutation as indicated. For Y2H studies, ADAR1 domains were incorporated into the prey pGBKT7 AD plasmid, including ADAR1 p150* FL (p150, aa1-1226), ADAR1 Zαβ (Zab, aa1-502), ADAR1 Zαβ W195A (Zab, aa1-502), ADAR1 Za (Zα, aa133-199), ADAR1 Zβ (Zb, 293-357), ADAR1 dsRBD1-3(RBD, aa459-aa795), ADAR1 dsRBD1-3 deficient mutant (RBDmut, aa459-aa795), and deaminase (Deam, aa795-1226) domains. In addition, OASL domains were cloned into the bait pGBKT7 BD plasmid, including OASL FL (OASL, aa1-514), N-OAS (aa1-349), Ubiquitin-like domains (UBLs, aa350-514), and UBL1 (aa354-433) and UBL2 (aa434-509). Coding sequences (CDS) of OASL and ADAR1 were subcloned into various backbone vectors as indicated, including pIRES -3XF-EF1α, pIRES-V5-EF1α, pCDNA3 HA-CMV, pmCherryC1-CMV, pEGFPC1-CMV, pCDH puro-CMV, and pCDH hygro-CMV. The HA-CMV vector was a gift from Edward Yeh (Addgene plasmid #18712). For *in vitro* protein interaction studies, the N-OAS construct (aa1-350) with an N-terminal GST tag was cloned into the PET28a vector. The ADAR1 dsRBD1-3 (RBD) construct (aa459-795) and ADAR1 Zα (Za) construct spanning aa133-199 with an N-terminal 6XHis tag were similarly cloned into the PET28a vector.

### Mass spectrometry

Immunoprecipitated products were separated by SDS-PAGE and fixed overnight prior to silver staining. Protein bands were excised and sent to the Taplin Biological Mass Spectrometry Facility (Harvard Medical School) for protein identification.

### Y2H screening

Complementary DNA encoding the human OASL full length (594 aa) was inserted into the bait vector pGBKT7. The AH109 yeast strain (*Saccharomyces cerevisiae*, Clontech), was transformed using two transformation rounds first with 0.5 µg of bait OASL and then 50 ug of cDNA prey human leukocyte cDNA library using the PEG/lithium acetate method as per the Matchmaker GAL4 Two-Hybrid System 3 protocol (Clontech). At least 10^6^ transformants were plated on both control (SD-Leu-Trp) and selective media (SD-Leu-Trp-His). For the yeast co-transformation assay, bait vectors containing various OASL constructs were transformed with the prey vector pACT2 carrying ADAR1 or indicated domains and screened on double (2DO, SD-Leu-Trp) and triple (3DO, SD-Leu-Trp-His) dropout media.

### Co-immunoprecipitation (IP) and GST pulldown

HEK293T cells were transfected with the specified DNA plasmids using polyethylenimine (PEI; Polysciences). After 48 hours, cells were harvested, rinsed with DPBS (Sigma), and lysed in 1% NP-40 buffer containing 50 mM Tris-HCl (pH 7.5, Invitrogen), and 150 mM NaCl (Sigma), supplemented with 1 mM EDTA, 10% glycerol, and cOmplete^™^, EDTA-free Protease Inhibitor Cocktail (Roche). Lysates were sonicated or subjected to three freeze-thaw cycles, then centrifuged at 16,000g for 10 min. Supernatants (whole cell extracts (WCEs)) were filtered through a 0.45 μm polyethersulfone membrane (Thermo Fisher) and precleared with Sepharose beads (GE) by rotating at 4°C for 1 h. For co-immunoprecipitation, WCEs were incubated with the appropriate primary antibodies ON at 4°C, followed by a 2 h incubation with protein A/G agarose beads (Thermo Fisher). For GST pulldown, WCEs were incubated with glutathione-Sepharose beads (GE) at 4°C for 1 h. Beads were washed five times with 1% NP-40 lysis buffer containing 150–500 mM NaCl. Finally, the beads were eluted in 2× Laemmli buffer (Sigma) by heating at 95°C for 5 min and analyzed by immunoblotting. For co-immunoprecipitation with IFNβ treatment, cells were treated with IFNβ (0.1 ug/ul) for 48 h post-treatment and harvested for subsequent immunoprecipitation.

### RNA Immunoprecipitation (RIP)

HEK293T ADAR1 KO cells were transfected with the indicated plasmids using PEI for 48 h. Cells were lysed with lysis buffer [50 mM tris (pH 7.5), 150 mM NaCl, 1 mM EDTA, 1% NP-40, cOmplete protease inhibitor (Roche), and SUPERase·In RNase Inhibitor (0.1 U/μl) (Invitrogen)]. Lysates were incubated overnight at 4°C with HA agarose beads (Pierce^TM^ Anti-HA Agarose ThermoFisher). The beads were washed gently four times with tris-buffered saline [50 mM tris (pH 7.5), 150 mM NaCl, and SUPERase·In RNase Inhibitor (0.05 U/μl) (Invitrogen)]. Bound proteins were eluted with 2× Laemmli Sample Buffer after boiling at 95°C for 10 min. Western blot was used to assess input and pulldown efficiency. RNAs bound to the agarose beads were extracted with TRIzol (Sigma) and purified following the manufacturer’s protocol and proceeded further with RT-qPCR.

### Immunoblotting

Protein lysates were separated on SDS–PAGE gels, transferred to PVDF membranes, and immunoblotted with primary antibodies overnight at 4°C or 2h at RT, followed by secondary antibody incubation for 1h at RT. A list of antibodies used can be found in Table 1.

### CRISPR/Cas9 Knockout

Single-guide RNAs (sgRNAs) targeting OASL were designed using the CRISPR design tool from Synthego (https://www.synthego.com/products/bioinformatics/crispr-design-tool) and are shown in Table S5. Briefly, HEK293T cells were subjected to CRISPR/Cas9-mediated gene knockout using preassembled ribonucleoprotein (RNP) complexes composed of Synthego sgRNAs and recombinant Cas9 protein. RNPs were formed by incubating sgRNAs with Cas9 at a 2:1 molar ratio and delivered into cells via electroporation using the Neon Transfection System under optimized conditions (1,200 V, 20 ms, 2 pulses). Post-electroporation, cells were cultured under standard conditions. Clonal populations were generated by limiting dilution, and successful knockouts were confirmed by Sanger sequencing and Western blot analysis following Sendai virus infection (SeV, 100 HA) for 24h.

### Confocal, fluorescence, and immunofluorescence imaging

HeLa, HEK293T, and A549 cells were plated in 12-well plates containing glass coverslips ( Fisherbrand) and incubated overnight at 37 °C. Cells were then transfected for 24h or infected with virus as indicated for 24h. Cells were then washed with Dulbecco’s Phosphate-Buffered Saline (DPBS) (Gibco) and fixed with 4% paraformaldehyde for 15 min at room temperature (RT). Cells were permeabilized using 0.1% Triton X-100 in DPBS for 10 min and blocked with 1% BSA in DPBS containing 0.1% saponin for 2h at RT. Immunostaining was performed with primary antibodies incubated overnight at 4 °C or for 2h at RT, followed by three Phosphate-Buffered Saline containing 0.05% of Tween 20 (PBS-T) washes and incubation with respective secondary antibodies (1:300-1:1000, Alexa Fluor 488/594). Coverslips were mounted with ProLong Gold Antifade Mountant containing DAPI (ThermoFisher). Imaging was conducted using a confocal microscope (Leica TCS-SP8-AOBS Confocal (inverted)), Keyence (BZ-X810), or ZOE (Bio-Rad Fluorescent Imager) at indicated magnification. Image analysis was performed using LAS X and ImageJ software.

### Recombinant protein expression and purification

Recombinant domains of OASL and ADAR1 were purified as previously described.^25,57^ Briefly, proteins were transformed into human codon optimized Rosetta (DE3) *E. coli* cells and grown at 37**°**C to an OD600 of 0.5-0.7 in LB broth with 50 ug/ml Kanamycin and 30 ug/ml chloramphenicol. The culture was cold-shocked for 30 min at 4°C prior to induction with IPTG (1 mM). Induction was carried out at 18°C overnight (ON) for most constructs, or at 37°C for 3h for ADAR1 Zα. The cells were harvested by centrifugation and resuspended in 10 ml per liter of culture in lysis buffer (50 mM KH_2_PO_4_, pH 6.0, 500 mM NaCl, 20 mM imidazole, 5 mM β-mercaptoethanol, 0.5 mM PMSF, and Protease Inhibitor Cocktail (Sigma-Aldrich). Lysates were sonicated and cleared by centrifugation at 18,000 g for 1 h at 4°C. Affinity purification was performed using the batch method, 2 ml of Co-NTA agarose (Qiagen) slurry per liter of culture supernatant, incubated for 1 h at 4°C with gentle rotating. For ADAR1 RBD purification, the lysate-bead mixture was equilibrated with equilibration buffer [50 mM Tris-HCl (pH 7.5), 200 mm KCl, 10 mM imidazole, 20% glycerol, 0.5 mM PMSF, and 0.50% Triton X-100] as reported.^25^ The beads were washed three times with 10× column volume (CV) of wash buffer [50 mM Tris-HCl (pH 7.5), 200 mM KCl, 80 mM imidazole, and 0.50% Triton-X]. Co-NTA-agarose-bound proteins were eluted with 10× CV of elution buffer [50 mM Tris-HCl (pH 7.5), 200 mm KCl, 250 mM imidazole, 20% glycerol, 0.5 mM PMSF, and 0.50% Triton X-100]. For OASL domain purification, similar procedures were followed using 500 nM NaCl instead of 200 nM KCl as reported. Eluted proteins were concentrated to 1 ml using Amicon centrifugal filter concentrator (30 kDa MWCO, Millipore Sigma), desalted and buffer-exchanged using a PD-10 desalting column (GE healthcare) with 50 mM Tris-HCl (pH 7.5), 20% glycerol, and 150 mM NaCl. For ADAR1 Zα, the eluate was further purified using cation exchange chromatography (HiTrap SP HP 5 ml column, Cytiva) with a NaCl gradient on an AKTA system. Protein aliquots were snap-frozen in 25% glycerol and stored at -80°C until further use.

### RNA electrophoretic mobility shift assay (REMSA) and EMSA

The single-strand FAM-RNA (IDT, 50 µM) were synthesized along with an unlabeled complement strand (IDT, 50 µM) and annealed to dsRNA (20 µM) using 5X annealing buffer (50 mM Tris-HCl, pH 8.0, 100 mM NaCl, 5 mM EDTA). Both strands were mixed and heated at 90°C for 1 min and slowly let cool to RT. The FAM duplex was stored frozen at -20°C until use. The FAM-labeled 38 bp RNA probe (5’-FAM/AAUUAACCAAGGAAAAUAACAAGGACAGGGACCAGGUU) and 71 bp RNA probe (5’-FAM/GGGCUAUUCGUUUUCCUAUUGAGCAUAGCCGCUUCUUCGGCUAUGCUCAAUACUUUCGGAAAAUGAACAGU) were incubated with proteins at indicated concentration in binding buffer (50 mM Tris, pH 6.5, 1 mM EDTA, 0.5 mM DTT, 100 mM KCl, 5% glycerol) at 37°C for 30 min. Samples were resolved on a 6% native TBE polyacrylamide gel in 0.5x TBE buffer system, imaged using a ChemiDoc system (BioRad), and analyzed with ImageLab Touch Software (BioRad). For the 6XHis GST UBLs EMSA assay, the protein was incubated using increasing concentrations of 6XHis ADAR1 Zα protein and binding buffer ( 50 mM Tris, pH 7.5, 5% glycerol) at 30 min in RT. Samples were resolved on 6% native Tris-glycine polyacrylamide gels using Tris-glycine running buffer and electrophoresed at 80 V for 4 h at 4°C.

### Quantitative RT-PCR (RT-qPCR)

Total RNA was isolated using either TRI reagent or the RNeasy Mini kit, treated with DNase I (Sigma) to remove genomic DNA, and then reverse-transcribed into cDNA using the iScript cDNA synthesis kit (Bio-Rad). Quantification of cDNA was performed using the SsoAdvanced Universal SYBR Green Supermix (ThermoFisher) on a CFX96 real-time PCR system (Bio-Rad). The gene-specific primers used for qRT–PCR are detailed in the Supplementary Table.

Complementary DNA (cDNA) was synthesized using the iScript cDNA synthesis Kit (Bio-Rad) and quantitative PCR (qPCR) was performed using the SsoAdvanced Universal SYBR Green Supermix (ThermoFisher). RNA input levels were normalized to GAPDH and ratio was taken compared to EV. For RNA IP analyses, the % input was taken as 2^-dCT^. Primer sequences used are listed in Table S5.

### RNA editing by Sanger sequencing

RNA editing by Sanger sequencing was assessed in HEK293T and A549 WT cells expressing the indicated constructs. The cDNA was amplified by PCR using the Phusion High-Fidelity DNA Polymerase (NEB) and then purified with the PCR Purification Kit (Qiagen). PCR products were Sanger sequenced and analyzed using SnapGene Viewer. RNA editing frequencies were determined by measuring the peak areas of adenosine and guanosine using BioEdit. All significantly and validated editing sites are listed in Table S3. Primer sequences can be found in Table S5.

### Dual-Glo assay

HEK293T cells were transfected in 12-well plates with a total of 2 µg of plasmid DNA, including the editing reporter and specified plasmids using PEI. After 48 hours, cells were lysed with 1X Passive Lysis Buffer (Promega) for 30 min RT. Luminescence from Firefly and Renilla luciferases was measured using the Dual-Glo Luciferase Assay System (Promega) on a Molecular Diagnostics plate reader. Relative Light Units (RLU) were calculated as the ratio of Firefly (FLUC) to Renilla (RLUC) luminescence normalized to EV control.

### Stable cell line establishment

Stable expression in HEK293T and A549 cells was established via lentiviral transduction followed by antibiotic selection and validation by immunoblotting (Figure 4A). HEK293T cells were plated in 6-well dishes and transfected with 0.4 μg each of pReV and pVSV, along with 0.8 μg of the pGag/Pol packaging plasmid and the indicated expression constructs using PEI according to the manufacturer’s instructions. Three days post-transfection, virus-containing supernatants were harvested, filtered, and stored at –80 °C until use. Target cells were infected with lentiviral supernatants in the presence of 8 μg/ml polybrene for 48 h, followed by antibiotic selection using puromycin and/or hygromycin depending on the resistance cassette encoded by the lentiviral construct. Selection was maintained until non-transduced control cells were eliminated and stable populations were established. Expression of OASL, p150*, and co-expression lines was subsequently confirmed by immunoblot analysis prior to downstream functional assays.

### Total RNA library preparation

Total RNA samples were first evaluated for quality using the Agilent 4200 TapeStation, which provided RNA integrity numbers (RIN), and quantified using the Qubit Fluorometer (ThermoFisher). Only samples meeting the required quality criteria advanced to library preparation, which was performed according to the manufacturer’s instructions (Illumina). In brief, ribosomal RNA was removed, and the remaining RNA was fragmented and denatured. cDNA synthesis was performed in two steps: first-strand followed by second-strand synthesis. Subsequent steps included end repair, A-tailing, and adaptor ligation. The resulting libraries were PCR-amplified, assessed for quality, quantified, and finally sequenced using the Illumina NovaSeq 6000 platform.

### RNA-seq data processing and quality control

Bulk RNA-seq Paired-end (2×150 bp) sequencing was done for 12 samples from HEK293T cell line expressing empty vector (EV), OASL, ADAR1p150* and both (OASL+ADARp150*) in triplicates using Illumina NovaSeq platform at the LRI Genomics core. Raw fastq reads from multiple lanes were concatenated and merged into forward (R1) and reverse (R2) fastq files utilizing fastp. Quality filtering was performed using fastp v0.23.2 (30423086), with phred quality filtering (≤ 20), allowed unqualified percent bases (30%), auto-detect adapter removal, and deduplication (to drop duplicated reads).

### RNA-seq data analysis and differential gene regulation

Quality-filtered reads from each sample were aligned to the reference human genome build *hg38* utilizing STAR mapper (v2.7.10a) (23104886). Further, mapped BAM files were subjected to duplicates removal and indexing with samtools (v1.16.1) (33590861). Quality checks were performed using FastQC (v0.11.9) and MultiQC (v1.21) (27312411). The processed BAM files were used for the gene-wise read counts utilizing feature counts from Rsubread v2.16.1 (24227677, 30783653) package using inbuilt hg38 NCBI RefSeq meta-feature annotation with duplicates were ignored. For all the ENTREZIDs, gene symbols were retrieved using org.Hs.eg.db (v3.15.0) annotation package. Finally, filtering lowly expressed genes, log-CPM TMM normalization and differentially expressed genes (DEGs) regulation analysis was performed using edgeR (v3.38.4) (19910308) and limma-voom (v3.52.4) (25605792, 24485249) packages with setting up group wise design and contrast matrix, voomWithQualityWeights normalization, eBayes function. Genes with logFC (log fold-change ∼ 0.5) and FDR (false discovery rate) ≤ 0.05 were counted as DEGs. Group-specific DEGs were further used for the gene ontology and pathway enrichment analysis implementing Enrichr (27141961) and IPA (Ingenuity Pathway Analysis) tools. EnhancedVolcano (v1.20.0) and ggplot2 packages are utilized to show regulatory plots. All DEGs are listed in Table S4. Primer sequences can be found in table S5.

### RNA editing alterations and regulations

Further, the effect of OASL mediated RNA editing alterations were explored. Briefly, STAR (23104886) aligned and samtools (33590861) filtered RNA-seq BAM files and hg38 human genome build were utilized for the identification of RNA editing events using REDItools (31996844, 23742983) with default parameters with excluding multi hits, consider mapping quality and remove substitutions in homo-polymeric regions and using known edit annotation. For this, known editing events annotation from the REDIportals (33104797, 39588754) were prepared having ∼16 million sites into GFF (gene feature file) and tabix indexing is done using tabix (v0.2.6) module (21208982). Further, filtering and selection of editing events were performed using editing coverage, frequency level and excluding multiple substitutions and merging to make frequency matrix. These editing events were further annotated using REDIportals known genomic annotations with SNPs, and repeats (33104797, 39588754). Added, sample-wise overall editing index was calculated, and genomic-region specific editing distribution has been analyzed. Later, significant dysregulated editing events were determined using Student’s t-test and Mann-Whitney (MW) U test was also performed (32211029), however it is restricted due to limited number of samples. RNA edited genes were subjected to pathway and gene ontology enrichment analysis using Enrichr (27141961), ShinyGO 0.85 (31882993), and IPA tools. Association and differential editing results were drawn as Manhattan and volcano plots implementing ggplot2 and CMplot packages.

### Virus Stock

Parental VERO/hSLAM cells (Sigma 04091501-1VL) were cultured in Dulbecco modified Eagle’s medium (DMEM) supplemented with 5% FBS, penicillin (100 μg/ml), streptomycin (100 units/ml) (Gibco/Invitrogen), and 0.4 mg/ml Geneticin (ThermoFisher). The recombinant Measles Virus Edmonston vaccine strain including a gene encoding mCherry fluorescent protein was provided by Dr. Jianrong Li (The Ohio State University). The MeV Schwarz strain was purchased from NISBC (Measles Vaccine (Live) 2nd International Reference Reagent 92/648). MeV strains were propagated by infecting VERO/hSLAM cells at MOIs of 0.2 TCID_50_/cell for 5-7 days at 32°C as previously described.^58^ The GFP VSV was generously obtained from Dr. Sean Whelan (Washington University). The Sendai virus Cantell strain was purchased from Charles River.

### Virus infection

A total of E6 A549 stably expressing cells were seeded in a 6-well plate. At ∼60% to 80% confluency, cells were washed with DPBS and media was changed to DMEM containing 2% FBS for virus infection. Cells were infected with in-house GFP-tagged VSV (MOI: 0.05), MeV Schwarz vaccine strain (MOI: 0.1), or mCherry-tagged MeV Edmonston vaccine strain (MOI: 0.01) and incubated at 37°C. For VSV, supernatants were collected at 0, 6, 12, 24, and 36 hpi. For MeV, samples were collected through multiple freeze-thaw sessions and combined with the extracellular virus in the supernatant at 0, 1, 2, 3, and 4 dpi. For rescued protein function in HEK293T DKO cells, cells were transfected for 24 h and then infected with VSV ( MOI: 0.001) or MeV (MOI: 0.005) at the designated time points.

### GFP VSV virus titration

Briefly, E6 VERO/hSLAM cells (Sigma 04091501-1VL) were seeded in a 12-well plate. Monolayers cultures were then infected with the A549 OE-generated VSV virus in 10-fold dilutions in triplicates using 2% FBS for 1h at 37°C while swirling. The virus inoculum was then replaced with 1% DMEM supplemented with sodium bicarbonate low melting point agarose in a 1:1 mixture overlay and let solidify for 15 min at RT. Plates were incubated at 37°C for 24h, fixed with 4% formaldehyde, and fluorescent plaques were counted using the Keyence microscope (BZ-X810) for titration. GFP+ colonies were counted as plaques for pfu/ml titration. Wells with at least five plaques were considered within acceptable limits. Statistical significance was determined using a two-way analysis of variance (ANOVA) followed by Tukey-Kramer post hoc multiple comparison tests, along with Student’s t-tests where applicable. Groups with a P value of less than 0.05 were considered significantly different.

### Construction of recombinant MeV Edmonston strain expressing mCherry

The mCherry gene was inserted into the junction between the P and M genes in the genome of measles virus (MeV) Edmonston strain using a yeast-based recombination technique as previously described.^59–61^ Briefly, six overlapping DNA fragments covering the entire genome of MeV were amplified by PCR. The mCherry gene, which contains the overlapping sequences with the P and M genes, was also amplified by PCR using two primers (mCherry-F: 5’ CGCCAGCCCATCAACGCGTATAATGGTGAGCAAGGGCGAGGAGGAT 3’; mCherry-R: 5’ TTTATAATGGATTTAGGTTGTATCATCATTTGTACAGCTCGTCCAT 3’). 200 ng of each MeV DNA fragment was mixed with 100 ng of pYES2 vector in PEG/LiAc solution, which were transformed into MaV 203 competent yeast cells by heat-shock and plated on SD/Ura-agar plates. Positive yeast colonies were initially screened by PCR. The positive plasmid from yeasts was transformed into *E. coli* DH10B competent cells, and plasmid DNA was verified by HindIII restriction enzyme digestion and sequenced to confirm that no additional mutations were introduced during the assembly. The full-length genomic cDNA of MeV containing mCherry was assembled into pYES2 vector under the control of T7 promoter and was flanked by a hepatitis delta virus ribozyme (HDVRz) sequence and a T7 terminator. The final plasmid was designated as pMeV-mCherry. Recovery of recombinant MeV (rMeV-mCherry) from the infectious clone was carried out as described previously.^59–61^ Briefly, pMeV-mCherry and three support plasmids encoding MeV ribonucleocapsid complex (pN, pP, and pL) were co-transfected into HEp-2 cells which were infected with a recombinant modified vaccinia Ankara virus (MVA-T7) expressing T7 RNA polymerase.^62^ Four days later, cells and supernatants were collected, and co-cultured with Vero CCL81 cells. Syncytia, typical cytopathic effects of MeV infection, was observed at day 2 after co-culture. The recovered rMeV-mCherry was further amplified in Vero CCL81 cells. Subsequently, the virus was plaque purified as described previously.^59^ Individual plaques were isolated, seed stocks were amplified in Vero CCL81 cells, and sequencing confirmed. The viral titer was determined by a plaque assay or TCID_50_ assay in Vero CCL81 cells.

### mCherry MeV Edmonston titration (FU and TCID_50_)

For the foci forming unit assay, 0.3E6 VERO/hSLAM cells were seeded in 12-well plates in 10% FBS DMEM media + 1% Penicillin/Streptomycin L-Glutamine for overnight incubation. At 100% confluency, cells were washed with DPBS prior to virus inoculation using 2% FBS DMEM. Cells were infected with 10-fold dilution of viruses with at least three replicates for 1h and the media was replaced with1% agarose overlay for 7 dpi at 32°C to maintain vaccine strain phenotype. Plates were then fixed with 4% formaldehyde and imaged using a Leica inverted microscope plate scanner (Imaging Core). A549-OE-generated mCherry MeV Edmonston titers were also assessed by measuring the TCID_50_ using an endpoint dilution assay with at least eight replicates per concentration following the Reed-Muench method (https://www.virosin.org/tcid50/TCID50.html). E6 cells/well of VERO/hSLAM cells were seeded in 96-well plates and monolayer of culture were inoculated with 10-fold virus dilution at 32°C for 7 dpi. Cells were washed with DPBS and fixed with 4% formaldehyde. Fluorescence was measured using a Leica inverted microscope plate scanner. Wells with mCherry fluorescence were counted as positive.

### Quantification and statistical analysis

All data were analyzed using Prism software (*v10.0.2, GraphPad*). Statistical analyses were performed using Student’s *t* test, one-way ANOVA, or two-way ANOVA as indicated. Unless otherwise indicated, data from *in vitro* editing assay and RNA-seq are expressed as mean ± SD from at least three independent biological replicates. For EMSA, data are presented as mean ± SD from at least two independent biological experiments. Differences not marked were considered not significant (n.s., *p* > 0.05). Statistically, the significance is indicated as ∗, *p* ≤ 0.05; ∗∗, *p* ≤ 0.01; ∗∗∗, *p* ≤ 0.001; ∗∗∗∗, *p* ≤ 0.0001. Details are provided in figure legends.

## SUPPLEMENTAL

Table S1. Mass Spectrometry Analysis of Endogenous OASL-Interacting Proteins.

Table S2. Yeast Two-Hybrid Screening Results.

Table S3. Differential RNA Editing Analysis.

Table S4. Differential Gene Expression Analysis.

Table S5. Primer Sequences.

